# Divergent recovery trajectories in reef-building corals following a decade of successive marine heatwaves

**DOI:** 10.1101/2023.07.16.549193

**Authors:** Kristen T. Brown, Elizabeth A. Lenz, Benjamin H. Glass, Elisa Kruse, Rayna McClintock, Crawford Drury, Craig E. Nelson, Hollie M. Putnam, Katie L. Barott

## Abstract

Increasingly frequent marine heatwaves are devastating coral reefs. Corals that survive these extreme heat stress events must rapidly recover if they are to withstand subsequent events, and long-term survival in the face of rising ocean temperatures may hinge on recovery capacity and acclimatory gains in heat tolerance over an individual’s lifespan. To better understand coral recovery trajectories in the face of successive marine heatwaves, we monitored the responses of bleaching-susceptible and bleaching-resistant individuals of two dominant coral species in Hawaiʻi, *Montipora capitata* and *Porites compressa*, over a decade that included three marine heatwaves. Bleaching-susceptible colonies of *P. compressa* exhibited beneficial acclimatization to heat stress (i.e., less bleaching) following repeat heatwaves, becoming indistinguishable from bleaching-resistant conspecifics during and after the third heatwave. In contrast, bleaching-susceptible *M. capitata* repeatedly bleached during all successive heatwaves and exhibited seasonal bleaching for up to three years following the third heatwave. Encouragingly, bleaching-resistant individuals of both species remained pigmented across the entire time series; however, pigmentation did not necessarily indicate physiological resilience. Specifically, *M. capitata* displayed incremental yet only partial recovery of symbiont density and tissue biomass across both bleaching phenotypes up to 35 months following the third heatwave. Conversely, *P. compressa* appeared to recover across most physiological metrics within two years, reverting to predictable seasonal variability. Ultimately, these results indicate that even some visually robust, bleaching-resistant corals can carry the cost of recurring heatwaves over multiple years, leading to divergent recovery trajectories that may erode coral reef resilience in the Anthropocene.

**Significance Statement:** Coral reefs are in jeopardy as climate change has led to increasingly frequent marine heatwaves. Some corals can survive these extreme heat stress events, thus acquiring environmental memory that may prime them for increased resistance and resilience in subsequent heatwaves via beneficial acclimatization. Yet, as the time between heatwaves decreases, the accumulation of stress experienced by some individuals may preclude opportunities for beneficial acclimatization. This nearly decade-long study revealed divergent recovery trajectories within and between species in response to successive marine heatwaves, ranging from costly to beneficial. As the climate continues to change, surviving corals must not only gain heat tolerance, but also rapidly recover to maintain the critically important ecosystem services that humanity relies on.

## Background

Ocean warming driven by climate change has led to staggering losses of live coral on coral reefs worldwide, and is among the most pressing of stressors threatening the survival of coral reefs today (1, 2). As mean ocean surface temperatures have steadily increased, there has been a corresponding increase in the occurrence of marine heatwaves (3, 4). These extreme warm water events can persist for days to months, frequently leading to coral bleaching, a symptom of the breakdown of the symbiosis between corals and their dinoflagellate algal endosymbionts (family Symbiodiniaceae) (5). This symbiosis is the energetic foundation of coral reef ecosystems, allowing for high rates of productivity in otherwise oligotrophic seas (6). The energy corals receive from symbiont photosynthesis supports a majority of their metabolic demand (7), thus fueling the construction of the complex three-dimensional framework necessary to support the most biodiverse ecosystems in the ocean (8). Given this significance, the breakdown of the coral-algal symbiosis during coral bleaching can have devastating consequences for coral reef ecosystems, ranging from declines in primary production to widespread coral mortality and reef erosion (1, 9). These losses can lead to concomitant declines in biodiversity and ecosystem function that harm not just these ecosystems, but also the human societies that rely on the myriad services functioning coral reefs provide (10). As marine heatwaves become increasingly frequent and severe (11), coral bleaching events are predicted to correspondingly increase (12). However, the extent to which individual corals can acclimatize to these conditions within their lifetime and thus gain resistance to recurring marine heatwaves remains a critical outstanding question.

Environmental memory of thermal stress, defined as the retention of information from an initial exposure that modifies the response to a later exposure, has been posited to lead to beneficial acclimatization of individuals exposed to repeat marine heatwaves (13–16). Indeed, coral communities that experienced sublethal heat stress and/or bleaching during marine heatwaves were less prone to bleaching in subsequent exposures across the Caribbean (17, 18), Great Barrier Reef (19), and Indo-Pacific (15, 16, 20, 21). There are likely multiple factors driving these increases in coral community heat tolerance, including beneficial acclimatization of surviving individuals (e.g., epigenetic modifications) (13–15), shifts in community composition driven by losses of bleaching-susceptible individuals (22–26), and selection for thermally tolerant offspring (i.e., adaptation) (27). However, exposure to sub-lethal heat stress in individual corals may not always lead to benefits in subsequent exposures (28), as a history of heat stress could exacerbate the effects of subsequent heat stress (i.e., sensitization) when corals have not fully recovered (29–31). In light of the increasing frequency and severity of marine heatwaves (11), there is a growing need to better understand how individual corals are responding to and recovering from repeat heatwaves and the ecological consequences of environmental memory for reef resilience into the future. Encouragingly, phenotypic diversity in bleaching susceptibility varies both within and between coral species, which is likely an important source of adaptive variation (32) that could lead to directional selection of bleaching resistance (e.g., (20)). However, the pace of ocean warming requires rapid acclimatization of established adult colonies in order to buy time for adaptation and proliferation of the next generation of thermally-tolerant corals, which may take several decades to centuries in these long-lived and slow-growing species (33). As marine heatwaves have only recently begun occurring on multi-decadal time-scales (1), we are just beginning to understand the capacity for individuals to recover, acclimatize, or sensitize following heat stress (14). Further, understanding how phenotypic variation in these responses between individuals and species influences population dynamics in the field remains underexplored, yet has important implications for predicting ecosystem diversity and function in the face of a rapidly changing climate (5).

In order to address this critical gap in our knowledge, we examined the responses of coral communities and individual colonies of two dominant reef-building coral species in Hawaiʻi, *Montipora capitata* and *Porites compressa,* following three marine heatwaves (2014, 2015 and 2019). Adjacent conspecific colonies with contrasting bleaching phenotypes (i.e., bleaching-resistant vs. bleaching-susceptible) were first identified during the second of two consecutive coral bleaching events that occurred in Kāne‘ohe Bay in 2014 and 2015 (34). Corals and reef-wide bleaching prevalence were tracked across the next nine years (2015–2023), which included another bleaching event in 2019 (35). Following the 2019 event, we repeatedly assessed a suite of physiological parameters (e.g., metabolic rates, symbiont densities, host biomass and tissue composition) over three years to better understand intra- and interspecific variation in recovery trajectories following repeat heat stress. In addition, we explored the capacity for environmental memory to influence coral thermotolerance through a standardized short-term acute heat stress assay (36). This investigation of individuals that encompass the phenotypic extremes within each population over nearly a decade in the field is a powerful framework to test the hypothesis that divergent individual responses to repeat *in situ* heat stress lead to variation in recovery trajectories, with consequences for future heat stress responses.

## Results

### Three successive heatwaves influenced reef-wide coral bleaching prevalence

Between 2014–2022, three significant thermal anomalies were documented across Kāne‘ohe Bay (Fig. 1, Figs. S1–S2, Tables S1–S2). During these events, maximum sustained degree heating weeks (DHW) at patch reef 13 (PR13) were lower in 2014 (7.3°C wk^-1^) than 2015 (8.8°C wk^-1^), and lowest in 2019 (5.1°C wk^-1^) (Fig. 1, Tables S1–S2). In contrast, 2019 was the most severe of the three events in the southern end of the bay and regionally (10–14°C wk^-1^; Table S2). From 2015–2023, coral cover at PR13 was dominated by *Porites compressa* (53.5% ± 0.9) and *Montipora capitata* (12.5% ± 0.5) (Fig. S3). At the peak of the 2015 heatwave, reef-wide bleaching prevalence reached 39.1% for *M. capitata* and 44.9% *P. compressa*, declining two months after the heatwave in *P. compressa* (1.4%), but not *M. capitata* (35.5%) (Fig. 2). The following summer, bleaching prevalence across PR13 reached 38.1% for *M. capitata* in the absence of any measurable heat stress, but was rarely observed for *P. compressa* (0.2%) (Fig. 2). This cyclic seasonal pattern of decreased bleaching prevalence of *M. capitata* in winter and increased bleaching prevalence in summer continued the following year. At the peak of the 2019 heatwave, reef-wide bleaching prevalence increased to similar levels observed in the 2015 heatwave in both *M. capitata* (45.9%) and *P. compressa* (38.8%) (Fig. 2); however, despite similar prevalence, the severity of bleaching was three-fold lower for *P. compressa* in 2019 than 2015 (35). Notably, bleaching prevalence again showed a cyclic seasonal pattern in *M. capitata*, declining in winter and increasing in each of the three summers following 2019 (September 2020: 46.9%; October 2021: 21.7%; September 2022: 14.9%). In contrast, reef-wide bleaching prevalence of *P. compressa* did not change between seasons and remained low (0.7–3.2%) (Fig. 2).

**Fig. 1.**
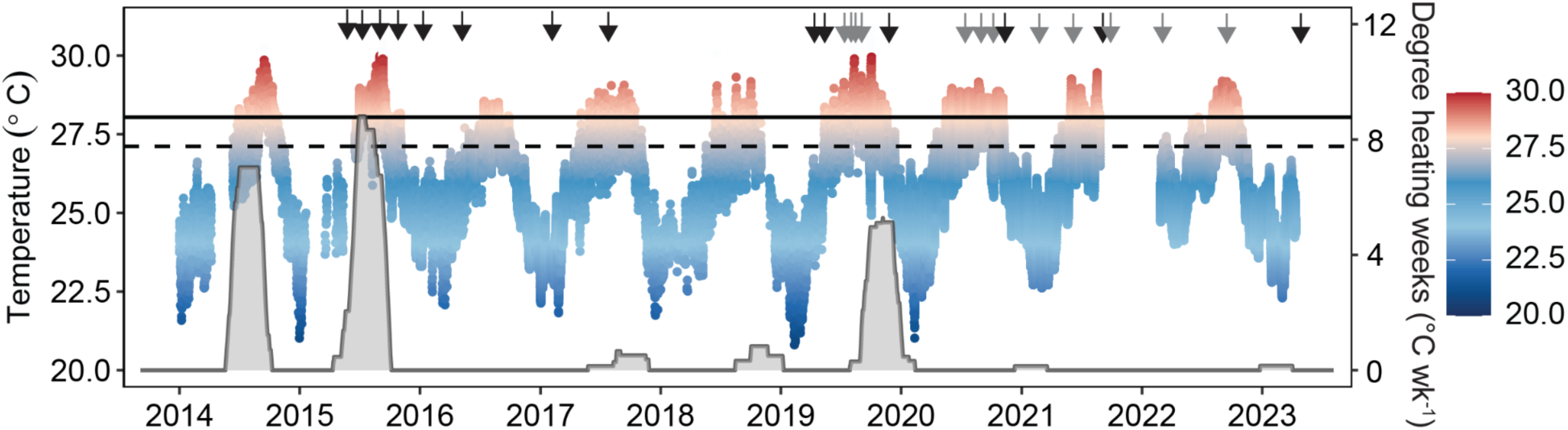
Temperature profile and heat stress accumulation over time. *In situ* temperatures were recorded from January 2014 – April 2023 at a depth of 0.7 – 2.7 m in Kāne‘ohe Bay. Points indicate hourly measurements (metadata in Table S1). Heat stress accumulation was estimated by degree heating weeks (gray shading) calculated from mean daily (24 hr) temperatures. Black arrows indicate image collection and gray arrows indicate image collection/physiological sampling. Dashed horizontal line indicates the Kāne‘ohe Bay’s climatological maximum monthly mean (MMM; 27.3°C) and solid horizontal line indicates the local coral bleaching threshold (MMM+1°C; 28.3°C).

**Fig. 2.**
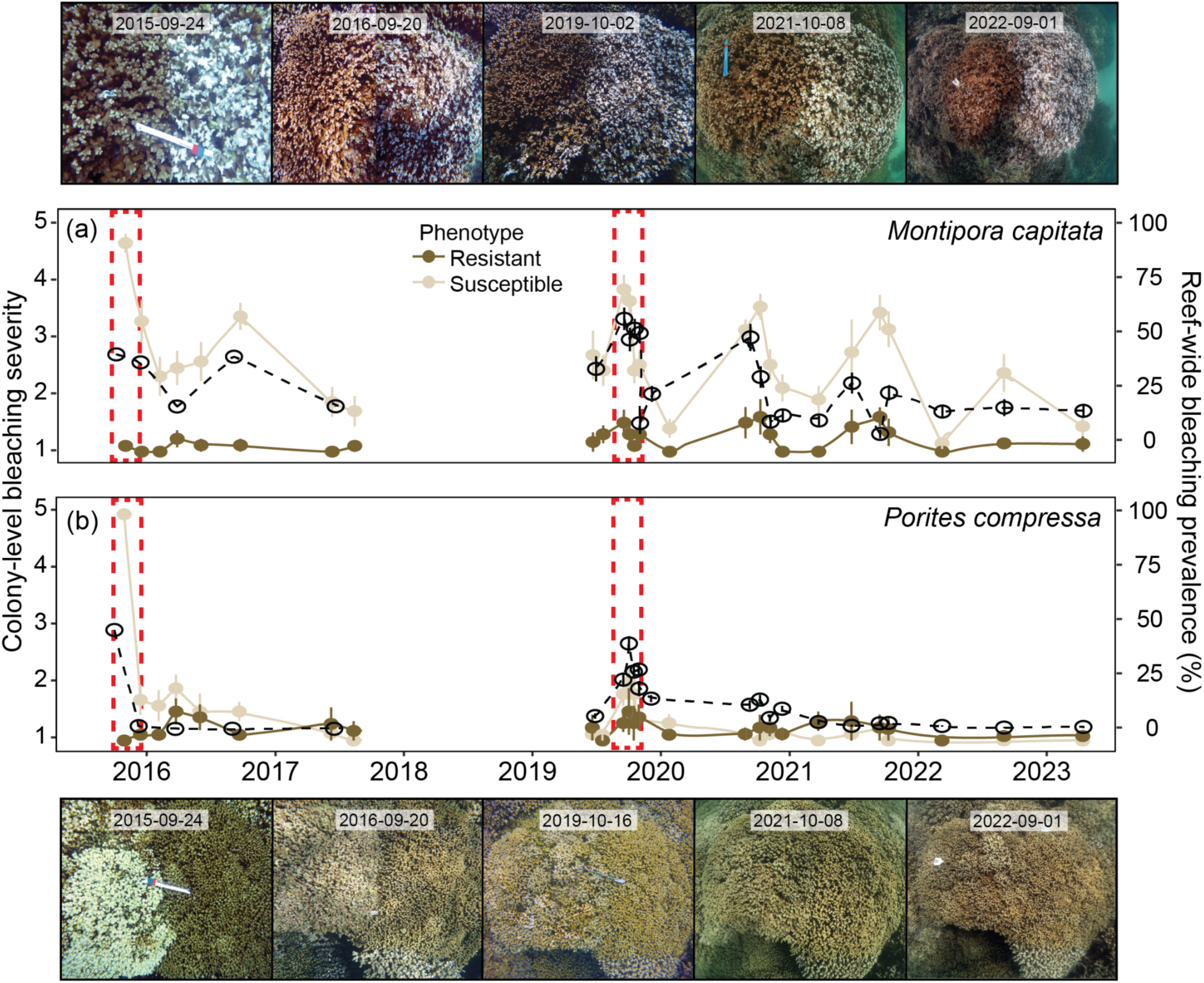
Bleaching response of bleaching-resistant and bleaching-susceptible corals over time. (a) Representative images of a *Montipora capitata* pair (left: bleaching-resistant, right: bleaching-susceptible) and mean colony-level bleaching severity (mean ± SE; n = 9–10) from September 2015 – September 2022. (b) Representative images of a *Porites compressa* pair (left: bleaching-susceptible, right: bleaching-resistant) and mean colony-level bleaching severity (n = 7–10). Colony-level bleaching severity was visually determined for each colony following the methodology of (35), where (1) represents 0% bleached or pale and (5) >80% bleached. Open circles and dashed lines represent mean reef-wide bleaching prevalence (n = 2–4 transects), calculated from benthic surveys for species of interest. Dashed red rectangles indicate the 2015 and 2019 marine heatwaves.

### Acclimatization and sensitization of individual colonies across recurring heatwaves

Colony-level bleaching severity was influenced by the significant four-way interaction between coral species, phenotype, season (i.e., winter v. summer), and time (F = 6.52, p < 0.0001). During the 2015 heatwave, bleaching-susceptible *M. capitata* and *P. compressa* were less pigmented than bleaching-resistant conspecifics (p < 0.0001) (Fig. 2). Throughout the next two years (2016–2017), bleaching-susceptible *M. capitata* remained less pigmented than bleaching-resistant conspecifics regardless of season (p ≤ 0.04) (Fig. 2). Interestingly, *P. compressa* phenotypes were indistinguishable by the summer after the heatwave (p = 0.23), and remained so throughout 2017 (Fig. 2). During the 2019 marine heatwave, bleaching-susceptible *P. compressa* (p < 0.03) and bleaching-susceptible *M. capitata* (p < 0.0001) were again less pigmented than bleaching-resistant conspecifics, although bleaching-susceptible individuals of both species were less severely bleached than the previous heatwave (Fig. 2). For the next two years (2020–2021), bleaching-susceptible *M. capitata* remained less pigmented than bleaching-resistant conspecifics regardless of season (p < 0.0001), whereas *P. compressa* phenotypes were indistinguishable by winter 2020 and remained so throughout 2021–2022 (p > 0.49) (Fig. 2). By winter 2022, *M. capitata* phenotypes were indistinguishable for the first time during the study period following the initial bleaching (p = 0.5), yet in the subsequent summer, bleaching-susceptible *M. capitata* were again less pigmented than bleaching-resistant phenotypes (p < 0.0001) (Fig. 2).

### Intra- and interspecific trajectories in physiological metrics indicate drawn-out and incomplete recovery

Coral physiological parameters were assessed from October 2019 (during the marine heatwave) to September 2022 (35 months post-heat stress) (Fig. 3, Fig. S4). Host tissue biomass (χ^2^ = 15.9, p < 0.03) and host lipid densities (χ^2^ = 25.2, p < 0.0003) were significantly influenced by the interaction between coral species and time (Fig. 3). *Porites compressa* had greater host tissue biomass than *M. capitata* across the time series (p < 0.003), with biomass converging three years after the 2019 marine heatwave (p = 0.24) (Fig. 3a, Fig. S4). For both *M. capitata* and *P. compressa*, tissue biomass and lipid densities displayed strong seasonality, where tissue parameters were greatest March–June prior to annual coral spawning (Fig. 3). During seasonal temperature maxima, tissue biomass for both coral species did not differ from the 2019 marine heatwave in the first summer following the heatwave (2020, p > 0.44), although biomass eventually increased in the second summer (2021, p < 0.0008) and third summer (2022, p < 0.0001) (Fig. S4).

**Fig. 3.**
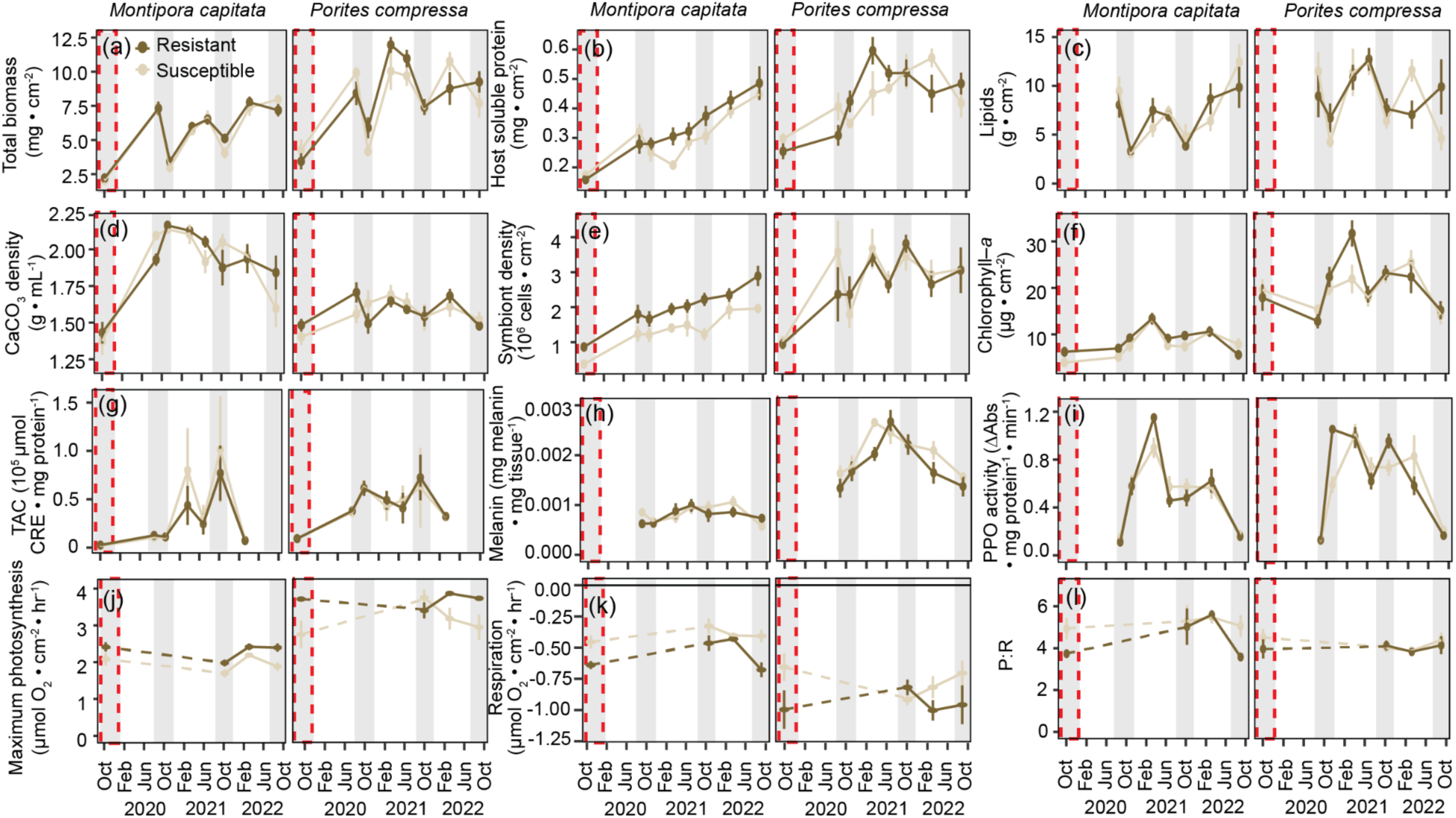
Physiological response of bleaching-resistant and bleaching-susceptible corals over time. (a) Host total tissue biomass (ash-free dry weight), (b) host soluble protein density, (c) host lipid density, (d) calcium carbonate (CaCO_3_) density, (e) endosymbiont cell density, (f) chlorophyll-*a* concentration, (g) host total antioxidant capacity (TAC), (h) host melanin content, (i) host prophenoloxidase (PPO) activity, (j) maximum photosynthetic rates, (k) light-enhanced dark respiration rates, and (l) photosynthesis to respiration ratios (P:R) for bleaching-susceptible and bleaching-resistant *Monitpora capitata* and *Porites compressa*. Grey shading indicates seasonal peak temperatures (i.e., September–October), with a dashed red rectangle representing the 2019 marine heatwave.

Host soluble protein density was significantly influenced by the interaction between coral species and time (χ^2^ = 20.5, p < 0.004). Generally, *P. compressa* had greater protein concentration than *M. capitata* across the time series (p < 0.008), apart from 10 months post-heat stress (p = 0.1), and later with protein concentration of the two species converging three years after the 2019 marine heatwave (2022; p = 0.55) (Fig. 3b). For *M. capitata*, protein concentration showed a stepwise increase over time, with concentration significantly greater across all timepoints when compared to the 2019 marine heatwave (p < 0.05). While bleaching-susceptible *M. capitata* protein concentration trended lower than bleaching-resistant conspecifics, there was no significant influence of phenotype (χ^2^ = 0.11, p = 0.74). For *P. compressa*, there was no difference in protein concentration between the 2019 marine heatwave and 10 months post-heat stress (p = 0.16), yet by 13 months, protein concentration was significantly greater across all subsequent timepoints (p < 0.01) (Fig. 3b, Fig. S4). Interestingly, during seasonal temperature maxima, *P. compressa* protein concentration was significantly lower 13 months post-heat stress than 24 months (p = 0.003), yet there were no differences between 24 and 35 months (p = 0.46) (Fig. S4).

Calcium carbonate (CaCO_3_) bulk density was significantly influenced by the three-way interaction between coral species, phenotype and time (χ^2^ = 14.1, p = 0.05). During the 2019 marine heatwave, CaCO_3_ density declined for both *M. capitata* and *P. compressa* regardless of phenotype, at which time density was indistinguishable between coral species (p > 0.72) (Fig. 3d, Fig. S4). Across the rest of the time series, CaCO_3_ density was significantly greater in *M. capitata* when compared to *P. compressa* regardless of phenotype (p < 0.05), with the exception of bleaching-susceptible corals at 35 months post-heat stress (p = 0.52). For bleaching-susceptible *M. capitata*, CaCO_3_ density was significantly greater across all timepoints when compared to the 2019 heatwave (p < 0.0001), again with the exception of 35 months post-heat stress when CaCO_3_ density was depressed (p = 0.59) (Fig. 3d). For bleaching-resistant *M. capitata*, however, CaCO_3_ density was significantly greater across all timepoints when compared to the 2019 heatwave (p < 0.002) (Fig. 3). Although bleaching-susceptible *P. compressa* trended lower than bleaching-resistant conspecifics during and 10 months after the marine heatwave, pairwise comparisons revealed that, for *P. compressa* of either phenotype, CaCO_3_ density was not significantly different across time (p > 0.05) (Fig. 3d, Fig. S4).

Symbiont densities were significantly influenced by the interaction between coral species and time (χ^2^ = 16.4, p = 0.02). While bleaching-susceptible *M. capitata* symbiont densities trended lower than bleaching-resistant conspecifics, there was no significant influence of phenotype (χ^2^ = 1.15, p = 0.28) (Fig. 3e, Fig. S4). Pairwise comparisons revealed that generally, *P. compressa* had greater symbiont densities than *M. capitata*, apart from during the 2019 heatwave when both species experienced declines (p = 0.25), and at 29 and 35 months post-heat stress (p > 0.07), when symbiont densities were greatest (Fig. 3e, Fig. S4). For *M. capitata*, symbiont densities did not rebound immediately after the 2019 heatwave, with symbiont densities indistinguishable from the 2019 heatwave until 17 months post-heat stress, after which symbiont densities increased over time. In contrast, *P. compressa* symbiont densities were significantly greater across all timepoints when compared to the 2019 marine heatwave (p < 0.02) and indicated seasonal patterns (Fig. 3e). For *P. compressa*, symbiont densities were significantly higher 24 months post-heat stress than 13 months (p = 0.0005), yet there was no difference between 24 and 35 months (p = 0.74) (Fig. S4).

Chlorophyll-*a* content was significantly influenced by the three-way interaction between coral species, phenotype and time (χ^2^ = 14.9, p = 0.04). Pairwise comparison revealed only one significant difference between bleaching-susceptible or bleaching-resistant phenotypes; that is, in March 2021, bleaching-susceptible *P. compressa* had lower chlorophyll-*a* content than bleaching-resistant conspecifics (Fig. 3f). The individual effects of time (χ^2^ = 24.67, p < 0.0008) and coral species (χ^2^ = 46.8, p < 0.0001) also emerged as significant, where *P. compressa* had significantly greater chlorophyll-*a* content than *M. capitata*. Post hoc analyses revealed for both species chlorophyll-*a* content did not differ from the 2019 marine heatwave until 17 months post-heat stress (March 2021) (p < 0.0001), when chlorophyll-*a* was greatest, yet, in June 2021 and September 2022, chlorophyll-*a* declined to values indistinguishable from the heatwave (p > 0.91) (Fig. 3f, Fig. S4). Metabolic rates, immunity (i.e., melanin content, prophenoloxidase) and total antioxidant capacity were influenced by the interaction of coral species and time, with all significant patterns detailed in the SI Appendix (Fig. 3g–l, Fig S4).

### Multivariate physiology reveals divergent responses across species and time

For both *M. capitata* and *P. compressa*, permutational multivariate analysis of variance (PERMANOVA) of physiological traits revealed a significant effect of time (p <0.0001). Pairwise comparisons revealed that multivariate physiology for all timepoints were significantly different from the marine heatwave for both species (p < 0.005) (Fig. 4). For *M. capitata*, all other comparisons were significantly different, apart from November 2020 and October 2021 (p = 0.11) and June 2021 and October 2021 (p = 0.21). For *P. compressa* after >1 year recovery from the marine heatwave, multivariate physiology indicated seasonal patterns, with no differences uncovered between August 2020 and September 2022 (p = 0.26), November 2020 and October 2021 (p = 0.16), and March 2021 and March 2022 (p = 0.62).

**Fig. 4.**
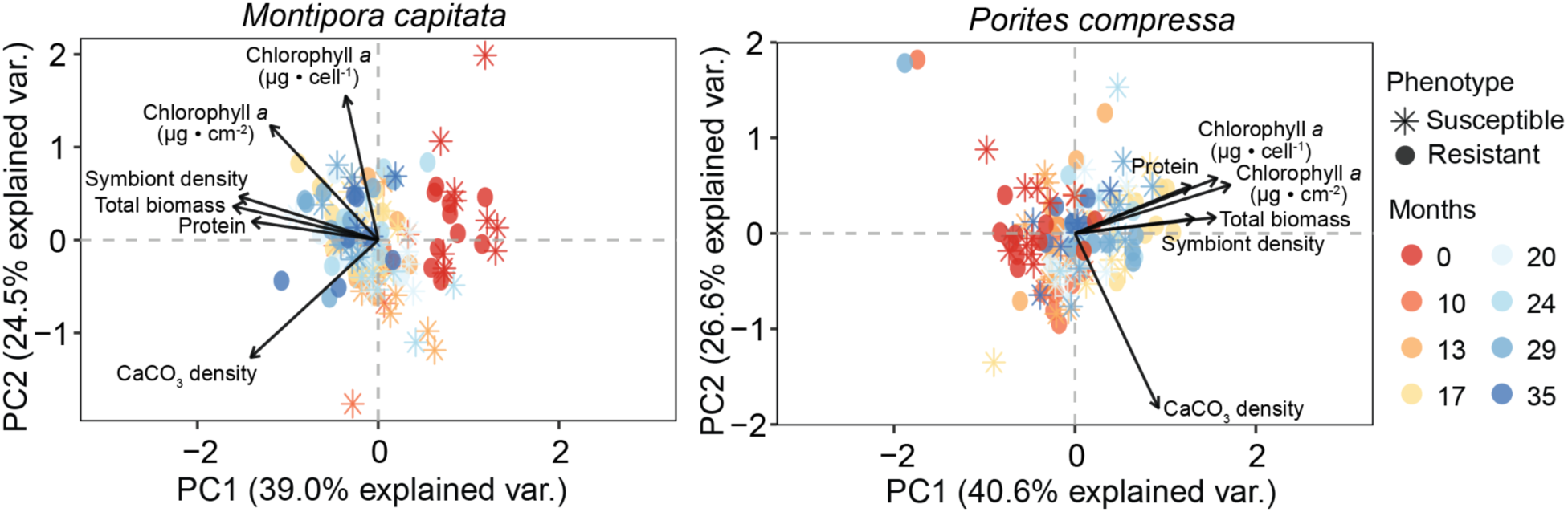
Principal components analysis (PCA) of physiological traits of bleaching-resistant and bleaching-susceptible coral phenotypes over time. Biplot vectors represent physiological traits (solid arrows), and points indicate the multivariate physiology of each coral genet (n = 7–10). Colors of the points indicate the months post-heat stress, where ‘0’ (red) represents the 2019 marine heatwave.

### Standardized acute heat stress assays confirm environmental memory influences heat tolerance

Photochemical yield (F_v_/F_m_) was significantly influenced by the interaction between treatment and species (Χ^2^ = 35.3, p < 0.0001) as well as treatment and phenotype (Χ^2^ = 9.1, p = 0.03). For both species, significant declines in photochemical yield were only observed in the most extreme treatment (p < 0.0001) (Fig. 5). Further, regardless of species, significantly lower photochemical yield was found in bleaching-susceptible compared to bleaching-resistant corals at +9°C (p = 0.01), whereas no differences in phenotype were found between the three other treatments (p > 0.74). There was a 1.18°C range in ED50 between the two species, with *P. compressa* more heat tolerant than *M. capitata*. Bleaching-susceptible *M. capitata* were the least heat tolerant, with a 50% reduction in F_v_/F_m_ observed at 36.1°C (95%CI: 35.8–36.5°C) vs. 36.6°C (95%CI: 36.3–36.9°C) for bleaching-resistant conspecifics (Fig. 5c). For *P. compressa*, bleaching-susceptible corals were virtually indistinguishable from bleaching-resistant phenotypes, with a 50% reduction in F_v_/F_m_ observed at 37.2°C (95%CI: 36.7–37.6°C) and 37.3°C (95%CI: 36.8–37.8°C), respectively. Color scores revealed similar patterns, with bleaching-susceptible *M. capitata* exhibiting lower ED50 (34.2°C; 95% CI: 33.3–35.1°C) than bleaching-resistant conspecifics (34.5°C; 95% CI: 33.7–35.3°C) (Fig. 5d). For *P. compressa*, color score ED50 was indistinguishable between bleaching-resistant and bleaching-susceptible *P. compressa* phenotypes (34.4°C and 34.6°C, respectively) (Fig. 5d).

**Fig. 5.**
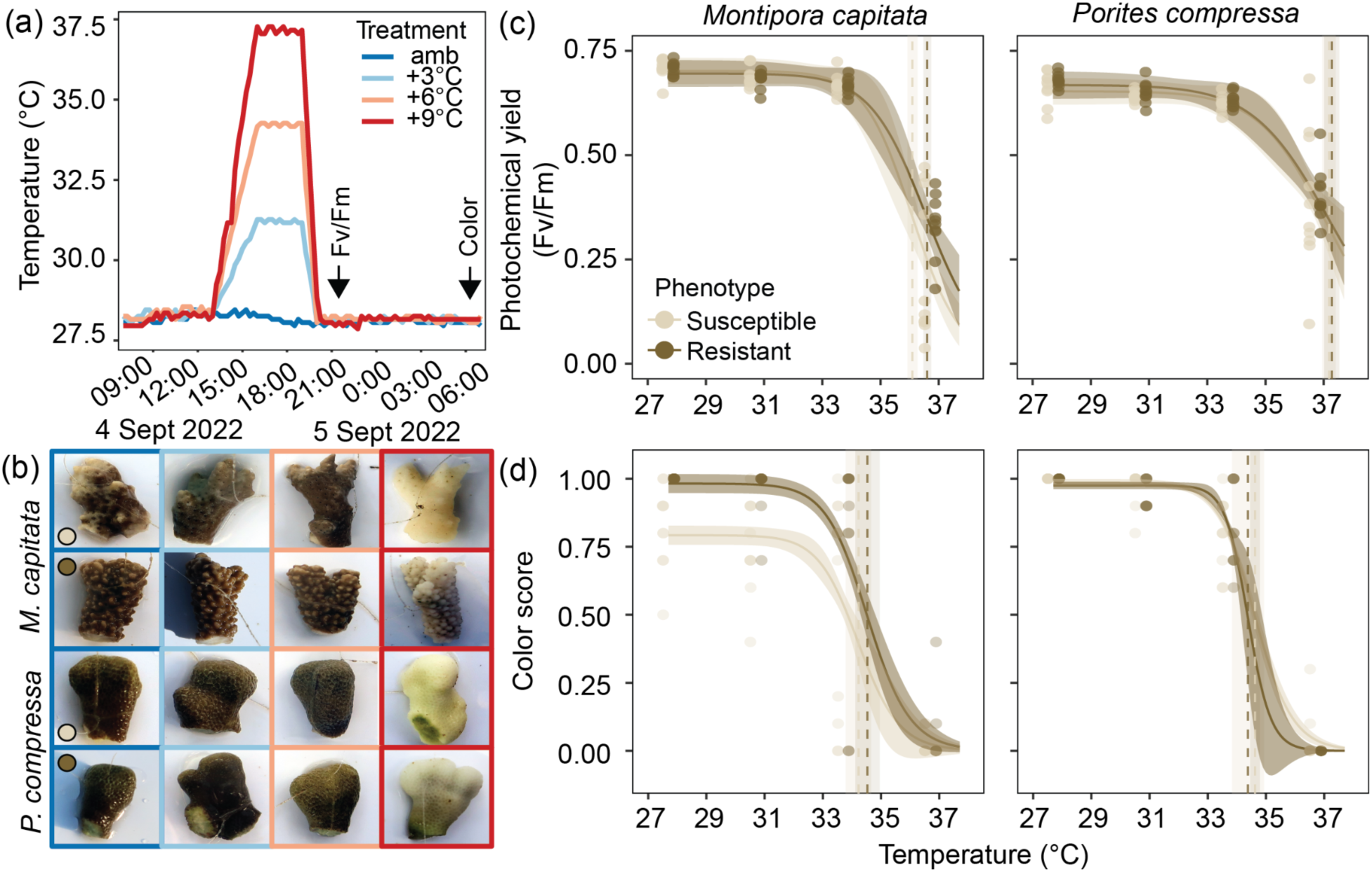
Variation in heat tolerance across bleaching-susceptible and bleaching-resistant corals from experimental heat stress assay. (a) Recorded hourly temperature measurements of the controlled experimental heat stress assays (amb = ambient temperature of 27.7°C), with an arrow indicating when dark-adapted photochemical yield (F_v_/F_m_) and color score were determined. (b) Respective images of coral fragments during the short-term heat stress. Three-parameter log-logistic dose–response curves were fitted to (c) F_v_/F_m_ measurements (± 95% CI) and (d) color score measurements (± 50% CI) in response to temperature for *Montipora capitata* and *Porites compressa*, where points indicate individual measures for coral genets (n = 10) in each treatment. Dashed vertical lines indicate the mean effective dose 50 (ED50) for each phenotype, with the standard error indicated by the shaded regions based on individual curve fits for each coral genet. Photochemical yield ED50s for each coral/phenotype were: 36.1°C *M. capitata* (susceptible), 36.6°C *M. capitata* (resistant), 37.2°C *P. compressa* (susceptible), 37.3°C *P. compressa* (resistant). Color score ED50s for each coral/phenotype were: 34.2°C *M. capitata* (susceptible), 34.5°C *M. capitata* (resistant), 34.6°C *P. compressa* (susceptible), 34.4°C *P. compressa* (resistant).

## Discussion

### Legacy effects of successive heatwaves have led to annual bleaching in susceptible *Montipora capitata*

Bleaching-susceptible colonies of *M. capitata* are now experiencing annual seasonal bleaching in the absence of anomalously high temperatures after a decade that included three marine heatwaves. This phenomenon was initially observed in the first summer following the 2015 heatwave, and was likely exacerbated by the combined impacts of the back-to-back heatwaves in 2014 and 2015 (37, 38). Encouragingly, in the second year after that heatwave, bleaching-susceptible *M. capitata* regained pigmentation over the winter and did not bleach again the following fall, indicating a ∼2-year recovery period. Yet, when faced with a third marine heatwave just four years later (2019), these same *M. capitata* colonies bleached again even though the heatwave was less severe. Declining performance in response to a second heatwave has been observed in corals from Australia (30) to the Caribbean (29); however, that bleaching-susceptible *M. capitata* colonies repeatedly bleached during each of the three summers following the 2019 event, even in the absence of thermal stress, is alarming. While seasonal declines in coral biomass and symbiont density during summer temperatures are common (39, 40), they do not typically lead to the visually apparent bleaching observed here. These results suggest that the frequency of heatwaves in Hawaiʻi over the past decade has compounded the stress experienced by susceptible corals, leading to persistent declines in performance under ambient conditions. More encouragingly, bleaching-resistant individuals within this same population have remained pigmented across multiple heatwaves, and were 0.5°C more heat tolerant than bleaching-susceptible conspecifics three years after the 2019 event. This suggests that when faced with future marine heatwaves, bleaching-resistant *M. capitata* will retain higher bleaching thresholds, which may promote survival (34) and favor the persistence of this species.

Despite the promise of reef resilience that bleaching-resistant *M. capitata* suggests, resistance to bleaching was not a sufficient measure of coral performance. Specifically, we observed incomplete recovery across multiple phenotypic and physiological metrics in both bleaching-resistant and bleaching-susceptible individuals in the three years following the 2019 marine heatwave, highlighting a legacy of stress that transcended visual bleaching assessments and persisted for several years. For example, neither phenotype exhibited complete recovery in symbiont or host protein densities during the three-year period following the 2019 heatwave. At their peak (after 35 months of recovery), symbiont densities only reached up to half of pre-heatwave densities (January 2014: 4–5 x 10^6^ cells cm^-2^ (41)). Given the ongoing upward trajectory, symbiont recovery will likely continue so long as another heatwave does not occur. These results underscore that physiological recovery can be a multi-year process, even when visual recovery is apparent within a few weeks to months following heat stress (38, 42). Interestingly, not all physiological parameters demonstrated a lag in recovery, with tissue biomass and lipid densities displaying apparent recovery followed by strong seasonality in the first year post-heat stress. However, other traits important for fitness, such as growth and fecundity, can take 4 years or more to recover following heat stress (18, 43–45). Long-term investigations are clearly needed to observe recovery in metrics closely linked to fitness as marine heatwaves continue to increase in frequency and severity, and thus increasingly overlap with the time required by many corals to fully recover. Decreasing periods of relief from thermal stress can compound physiological stress in many surviving corals, which can alter the relative performance within communities (31) and will have important ramifications for community composition and ecosystem function.

A better understanding of the mechanisms driving bleaching resistance may help predict coral fates in future reefs. For example, thermal tolerance in *M. capitata* is associated with a combination of host (32) and symbiotic (46) factors. In Kāne‘ohe Bay, *M. capitata* can host *Cladocopium* spp. and *Durusdinium glynnii* (47–49), and bleaching resistance is strongly correlated with higher proportions of *D. glynnii* (46). Indeed, the bleaching-susceptible colonies observed here hosted almost exclusively *Cladocopium* spp., while bleaching-resistant colonies predominantly hosted *D. glynnii* (32, 50) (Table S3). Both phenotypes exhibited additional physiological signatures of *Cladocopium*- or *Durusdinium*-dominated symbioses, respectively. For example, bleaching-susceptible *M. capitata* had lower symbiont densities than bleaching-resistant colonies across all seasons, matching others observations that *Cladocopium*-dominated *M. capitata* have lower symbiont densities than *Durusdinium*-dominated colonies (35, 38). While *Durusdinium*-dominated *M. capitata* were able to maintain bleaching resistance and higher symbiont densities, *D. glynnii* tends to provide the host with fewer resources than *Cladocopium* spp. under both ambient and heat stress conditions (51), indicating that symbiont retention during heat stress is an incomplete measure of coral performance. However, the nutritional benefits of hosting *Cladocopium* spp. may diminish over time as heatwaves and bleaching become increasingly common, and the impacts of changing symbiont dominance on coral calcification and thus ecosystem function require further study.

### Environmental memory of heatwaves has led to beneficial acclimatization in *Porites compressa*

Bleaching-susceptible and bleaching-resistant *P. compressa* appear to have converged on the same resistant phenotype despite past differences in bleaching susceptibility. Surprisingly, neither phenotype bleached during the 2019 heatwave, despite the bleaching-susceptible colonies exhibiting severe bleaching during the 2015 heatwave (34, 35). While lower bleaching severity may have been due to somewhat lower levels of heat accumulation in 2019, experimental heat stress tests confirmed identical bleaching thresholds in bleaching-susceptible and bleaching-resistant phenotypes. Together, these results suggest that beneficial acclimatization occurred and has persisted for several years. Higher bleaching thresholds have been observed in multiple coral species following successive marine heatwaves (17, 18, 20–22), suggesting that environmental memory of a prior stress event that improves the response to a subsequent exposure is common in corals. Increases in bleaching resistance from 2015 to 2019 were also observed in *P. compressa* across the population, which may also have stemmed from beneficial acclimatization; however, selection against weak corals or symbioses (20, 24, 31) or a shift to more stress-tolerant symbionts (31) is also possible. Support for these latter two hypotheses is limited for *P. compressa* in Kāne‘ohe Bay, where: (i) coral mortality was minimal in the aftermath of the 2014 heatwave (∼6%; (52)), (ii) *P. compressa* does not exhibit evidence of cryptic host speciation (53), and (iii) *P. compressa* maintains a specific symbiosis with a single symbiont species, *Cladocopium* ‘C15’ (47). Together, this suggests that the heatwaves in 2014 and 2015 resulted in stress hardening that first manifested as a decrease in bleaching severity in 2019, and this benefit has persisted for nearly a decade. While the exact mechanisms conferring this resilience have not been determined, physiological plasticity (18, 38), constitutive upregulation of stress-response genes (54, 55), and epigenetic modifications (56) all likely contribute, and represent important avenues of future study. Importantly, whether the benefits of environmental memory of moderate heat stress will persist as heatwaves become more intense remains unknown.

In the years following the 2019 heatwave, both phenotypes of *P. compressa* exhibited increases across multiple physiological traits, requiring ∼1.5 years to reach symbiont, chlorophyll-*a* and protein densities similar to historical pre-heatwave levels (57). Given their faster recovery and elevated bleaching thresholds (+0.7 – 1.2°C) relative to *M. capitata*, *P. compressa* may become more dominant in this region as heatwaves become more frequent. Indeed, *M. capitata* exhibited greater partial mortality following the 2015 event (34) and greater declines in benthic cover than *P. compressa* following the 2019 event (58). This likely favors the dominance of *P. compressa,* which already reaches >75% cover on some reefs in Kāne‘ohe Bay (59). While high coral cover has historically been considered a key metric of reef condition, the loss of biodiversity associated with transitioning from a multispecies assemblage to a predominantly *P. compressa* landscape will likely have a multitude of adverse effects on ecosystem function, from declining coral productivity to losses of blue food security (10, 60). Further, critical processes such reef accretion are projected to become uncoupled from coral cover under global change (61). Here, *P. compressa* was not able to sustain as high skeletal density as *M. capitata*, despite maintaining up to three times higher coral cover and exhibiting less bleaching. These patterns may be explained by: (i) enhanced heat tolerance resulting in trade-offs with growth ((62, 63) but see (64)), and/or (ii) the duration of our study was not long enough to capture the recovery of secondary calcification (i.e., densification), as recurring marine heatwaves could have delayed the revival of densification. Indeed, *P. compressa* are unable to recover calcification rates in the first eight months following heat stress (65), and elsewhere in the Pacific, *Porites* spp. can exhibit growth hiatuses for up to four years in the aftermath of a marine heatwave (45). Whether modern reefs are able to maintain net accretion and continue to provide the critical ecosystem services humans rely on remains to be seen.

## Conclusions

We have entered a new era of ocean warming that is affecting coral reefs in ways we are only just beginning to understand. Corals in the Main Hawaiian Islands have experienced a rise in offshore sea-surface temperatures of 1.15°C in the last 60 years (66, 67), leading to an unprecedented three successive coral bleaching events in the last decade (2014–2023) that were preceded by only a single widespread coral bleaching event the entire century prior (59, 68). The responses of corals to these increasingly frequent heatwaves demonstrates the divergent intra- and interspecific bleaching and recovery trajectories possible within a single coral community, highlighting the challenge of predicting future coral performance in a changing ocean. In one direction, *P. compressa* that were highly sensitive to marine heatwaves in 2014 and 2015 (i.e., severely bleached) have become visually and physiologically indistinguishable from bleaching-resistant conspecifics during a third heatwave (2019) and during extreme heat stress tests, suggesting beneficial acclimatization that persists across many years. This increase in bleaching resistance was accompanied by rapid physiological recovery after repeat heat stress, although there remained evidence of persistent stress (i.e., weakened skeletons), suggesting that these corals may undergo tradeoffs that can erode ecosystem function. Increases in coral bleaching resistance across recurring heatwaves has become more prevalent on reefs across the globe (15, 17, 19, 21), which is encouraging, as avoiding bleaching is associated with greater survival (34) and stress hardening may thus promote the persistence of corals in our warming oceans. On the other hand, bleaching-susceptible *M. capitata* are visibly struggling from the recent barrage of heatwaves, manifesting as seasonal bleaching in the absence of measurable heat stress. Importantly, even bleaching resistance was not associated with recovery capacity in *M. capitata*, highlighting the danger of predicting future individual performance and reef function from a lack of visual bleaching alone. The inability of *M. capitata* of either bleaching phenotype to recover physiologically after three years following a repeat heatwave underscores that coral resilience is a multi-faceted metric beyond bleaching resistance (62) and brings into question how we define coral resilience in the context of global change. Now more than ever, seasonal and long-term studies are critically needed to identify corals that cannot just withstand and survive repeated heat stress events, but also rapidly recover ecosystem-defining traits (e.g., biomineralization) to continue providing the critical ecosystem services coastal communities directly rely on. Urgent, collective global action to eliminate greenhouse gas emissions remains the only approach that may provide sufficient time for corals to acclimatize and adapt to rapid climate-induced temperature increases in order for coral reef ecosystems to persist in the Anthropocene.

## Materials and Methods

### Study site and seawater temperature

This study was conducted at patch reef13 (PR13) in the southern region of Kāneʻohe Bay, Oʻahu, Hawaiʻi. Seawater temperatures were recorded from January 2014 to April 2023. Where the record was incomplete, temperature data from two nearby reefs (<0.5 km away) were included. Cumulative heat stress (degree heating weeks; DHW) was determined following the equations in (69) using the maximum monthly mean (MMM) of 27.3°C. These results were compared to DHW on the reef of Moku o Loʻe (PR1), and versus DHW using the regional MMM of 27.0°C and MMM used in the literature for Kāneʻohe Bay (27.7°C (38); 28.0°C (35, 70)) (SI Appendix).

### Coral bleaching assessments

Adjacent conspecific corals with contrasting bleaching phenotypes (i.e., bleaching-resistant vs. bleaching-susceptible) were first identified during the 2015 marine heatwave (34). A total of 20 colonies each of *Montipora capitata* and *Porites compressa* (n = 10 colonies per phenotype per species) were followed for this study. Bleaching severity of each colony was determined from photographs taken between October 2015 to April 2023 (SI Appendix). Coral cover and reef-wide bleaching prevalence were also determined at each time point from benthic photoquadrats as in (34, 35) (SI Appendix).

### Physiological analyses

Fragments from each colony were collected across eight timepoints from October 2019 to September 2022. Photosynthesis rates were assessed at increasing irradiance via changes in oxygen evolution as previously described (35) (Fig. S5). Coral fragments were then flash frozen in liquid nitrogen and stored at −80°C until further processing. Coral tissue was removed from skeletons with a waterpik, and host and symbiont fractions were separated by differential centrifugation. All details of host (tissue biomass, protein and lipid concentrations, total antioxidant capacity, melanin content, prophenoloxidase activity, CaCO_3_ density) and symbiont (symbiont density, chlorophyll-*a* concentration) analyses can be found in the SI Appendix.

### Acute heat stress experiment

Ten individual colonies of each phenotype per species were sampled at the end of summer (September 1, 2022) and assessed for heat tolerance following a standardized protocol (36, 72). Briefly, corals were exposed to a 3-hour ramp to respective treatment temperatures (ambient: 28.0°C; amb+3°C: 31.0°C, amb+6°C: 34.0°C, and amb+9°C: 37.0°C), a 3-hour hold, and a 1-hour ramp down to ambient. At the end of the ramp and 1 h after sunset (∼19:30), corals were assessed for photochemical yield (F_v_/F_m_). The following morning corals were photographed with a color standard to assess coral color (Fig. S6, SI Appendix).

### Statistical analyses

All statistical analyses were done using R version 4.0.3 software (73), and are explained in detail in the SI Appendix. Briefly, a linear model was used to test for differences in colony-level bleaching severity between species, phenotypes, seasons, and time. Linear mixed effects models were used to assess for differences in physiological parameters. Differences in coral multivariate phenotypes were analyzed separately for each coral species using permutational multivariate analysis of variance (PERMANOVA) and principal components analysis (PCA), with the fixed effects phenotype and months post-heat stress. To determine how heat tolerance differed amongst coral species and phenotype, dose-response curves were fit to the median F_v_/F_m_ across temperature treatments and the effective temperature to induce a 50% loss in F_v_/F_m_ (effective dose 50; ED50) was calculated as the F_v_/F_m_ or color score from the model fit that is 50% of the initial value (72).

## Acknowledgements

Thank you to the staff of the Hawaiʻi Institute of Marine Biology for logistical support, especially marine safety officer Jason Jones. We also thank Chris Suchocki, Ariana Huffmyer, Wesley Sparagon, Mariana Rocha de Souza, and Joshua Hancock for field assistance. Corals were collected under the following permits from the State of Hawaiʻi’s Division of Aquatic Resources: SAP-2022-28, SAP-2021-41, and SAP-2020-41.

## Author contributions

Conceptualization: K.T.B, H.M.P., K.L.B.

Methodology: K.T.B, E.K., B.H.G., C.E.N., H.M.P., K.L.B.

Investigation: K.T.B, E.K., B.H.G., R.M., K.L.B.

Formal analysis: K.T.B.

Visualization: K.T.B, C.E.N.

Resources: E.A.L., C.D., C.E.N., H.M.P., K.L.B.,

Project administration: K.T.B, C.E.N., H.M.P., K.L.B.

Funding acquisition: C.E.N., H.M.P., K.L.B.

Writing - original draft: K.T.B, K.L.B.

Writing - review & editing: K.T.B, E.A.L., B.H.G., C.D., C.E.N., H.M.P., K.L.B.

