## Supplemental materials for "Divergent recovery trajectories in reef-building corals following a decade of successive marine heatwaves"

### Methods

*Study site and temperature data*

This study was conducted at patch reef 13 (PR13) in the southern end of Kāneʻohe Bay, Oʻahu, Hawaiʻi (21.4509, -157.7954). Hourly seawater temperatures were recorded continuously from January 2014 to April 2023 from temperature sensors within the reef at PR13 (1–2.7 m depth) as well as at two adjacent locations within 0.5 km: (1) PR12 (21.45096,-157.7972; 1.5 m depth) and (2) the National Oceanic and Atmospheric Administration (NOAA) Pacific Marine Environmental Laboratory (PMEL) 'CRIMP2' buoy (21.458, -157.798; 0.7 m depth) (Fig. 1, Fig. S1, Table S1). Mean daily (24 hours) seawater temperatures were calculated and averaged when data sources were overlapping, and used to determine cumulative heat stress (degree heating weeks; DHW) at PR13 following the equations in (Kristen T. Brown et al. 2022) (Fig. 1, Fig. S2, Table S2). The climatic maximum monthly mean (MMM) for Kāneʻohe Bay was determined from seawater temperature data from two monitoring stations on the reefs surrounding the island of Moku o Loʻe (PR1), located approximately 2 km from the study site in the southern region of Kāneʻohe Bay: 1) NOAA's MOKH1 Station[1] (21.433, -157.790; 1.7 m depth) and 2) the HIMB Point Lab Weather Station[2] (21.433, -157.7863; 1 m depth; (Rodgers, K. S., P. L. Jokiel, and Western Weather Group, Inc. 2005)). Data from 1992–2002 (excluding the 1996 marine heatwave) provided the closest available 10-year data to the time period used by NOAA for determining climatology MMM (1985–1990, 1993), resulting in a climatology MMM for Kāneʻohe Bay of 27.3°C. This MMM was used here to calculate DHW from 2014–2023. Cumulative heat stress at PR13 was compared to PR1 (21.4438, -157.7883; 1 m depth) (Table S1, Fig. S1). DHWs were also calculated from the temperature data recorded at both PR1 and PR13 using the MMM of 27.0°C (Main Hawaiian Island MMM as determined by NOAA and Kāneʻohe Bay climatology MMM from the 1960s (Jokiel and Coles 1977)), and other recent MMM values used for Kāneʻohe Bay in the literature: 27.7°C (Wall et al. 2021; Jury and Toonen 2019) and 28.0°C (Innis et al. 2021; Jury and Toonen 2019) (Fig. S2).

*Coral bleaching assessments*

Colony-level bleaching severity was determined from photographs of each colony following the methodology of (Innis et al. 2021), in which colonies were scored as: (1) no signs of paling (0%),

[1] https://www.ndbc.noaa.gov/station_page.php?station=mokh1
[2] http://www.pacioos.hawaii.edu/weather/obs-mokuoloe/#about

(2) mild paling (>20%), (3) moderate paling (20–50%), (4) mostly bleached (50–80%), and (5) fully bleached (80–100%). Observations occurred during peak and off-peak seasonal temperatures in most years. Benthic community composition was determined at the same time as colony-level observations following the same methods as in (Innis et al. 2021; Matsuda et al. 2020). Specifically, benthic photoquadrats (0.33 $m^2$), were imaged at 2 m intervals along a 40 m transect tape laid parallel to the reef crest at 1 m and 3 m depths (n = 1–2 per depth) at PR13. Benthic community composition was determined from each image via CoralNet using 50 randomly allocated points per photograph (Beijbom et al. 2015). Bleaching severity of each coral point was scored as: (1) pigmented (no signs of bleaching), (2) pale (moderately bleached), or (3) severely bleached (white). Reef-wide bleaching prevalence for each species was determined as the proportion of observations of that species showing signs of moderate or severe bleaching (i.e. bleaching score of 2 or 3).

*Physiological analyses*

A total of 276 fragments were collected from colonies of bleaching-resistant and bleaching-susceptible *M. capitata* and *P. compressa* (n= 7–10 pairs per phenotype per species) across eight timepoints from October 2019 to September 2022, incorporating the peak of the 2019 heatwave and three years (35 months) of recovery during non-heatwave years (Fig. S1). Fragments (4–5 cm in length) were collected by hand using bone cutters, transported to the Hawai'i Institute of Marine Biology (HIMB) in ambient seawater and held in flow-through seawater aquaria for 24–72 hours until measurements of coral performance. For four timepoints (October 2019, October 2021, March 2022, and September 2022), metabolic rates were assessed via changes in oxygen evolution using oxygen optodes (PSt7, PreSens) connected to an optical analyser (OXY-10, PreSens) (Innis et al. 2021). Oxygen optodes were calibrated on each day with a 0% oxygen solution (0.01 g $ml^{-1}$ $NaSO_3$) and 100% air saturated seawater. Coral fragments were analyzed between 08:00 and 17:00 within 250 ml clear acrylic chambers on top of a magnetic stirrer to allow for continuous mixing. Seawater temperatures were replicated to those experienced on the reef by using ambient seawater and a water jacket to maintain temperatures within the incubation chambers. Temperature and dissolved oxygen concentrations were recorded every 3 seconds at increasing increments of light over a total of 45–50 minutes to determine maximum net photosynthesis (Pmax). Light levels ranged from 112–726 μmol $m^{-2}$ $sec^{-1}$ in October 2019 (Innis et al. 2021) (step size ~100 μmol $m^{-2}$ $sec^{-1}$; ~10 minutes $step^{-1}$), and from 387–1800 μmol $m^{-2}$ $sec^{-1}$ in 2020–2021 (step size ~300 μmol $m^{-2}$ $sec^{-1}$; ~10 minutes $step^{-1}$). After measurements were completed at the maximum light levels, the lights were turned off (0 μmol $m^{-2}$ $sec^{-1}$) to measure light-enhanced dark respiration (LEDR) as defined by (Edmunds and Davies 1988). Photosynthesis-irradiance curves were fitted using the Platt model to extract Pmax and LEDR (Platt, Gallegos, and Harrison 1981; Innis et al. 2021). Upon completion of these *in vivo* analyses, coral fragments were flash frozen in liquid nitrogen and stored at −80°C until further processing.

Coral tissues containing symbionts were removed from skeletons with a waterpik using 50 mL of 0.1 M phosphate buffered saline (PBS) solution. The resulting holobiont homogenate was centrifuged at 4°C for 5 minutes at 2500 x g to separate the coral host fraction (supernatant) from the intracellular endosymbiont cells (family Symbiodiniaceae; pellet). Symbiodiniaceae were resuspended in PBS and cell abundances were quantified using a Millipore Guava flow-cytometer (Guava easyCyte 5HT) following the methodology of (Innis et al. 2021). Symbiodiniaceae cells were excited with a blue laser (488 nm) and identified by analyzing forward scatter and red autofluorescence in GuavaSoft 3.4 with the same gating for all samples. Symbiont densities were standardized to skeletal surface area ($cm^2$), which was determined using the wax-dipping technique (Holmes 2008; Stimson and Kinzie 1991). Endosymbiont photopigments were extracted in 100% acetone for 24 hours and the concentration of chlorophyll-*a* was determined via absorbance at 630, 663, and 750 nm using the equation in (Jeffrey and Humphrey 1975) and subtracting the absorbance at 750 nm from both A663 and A630:

$$Chlorophyll\ a = (11.43 \times (A663 - A750)) - (0.64 \times (A630 - A750))$$

Photopigment concentrations were standardized to both skeletal surface area and symbiont densities. Endosymbiont community composition (proportion *Durusdinium* vs. *Cladocopium*) of *M. capitata* colonies was determined by Dilworth et al. 2021 from samples collected in July 2019 using quantitative PCR.

Host tissue biomass was determined as ash-free dry weight (AFDW) from the coral fraction (i.e. symbionts removed as described above). First, 1 mL of the coral fraction was dried at 60°C for 24 hours until a constant weight was achieved. After the dry weight was recorded, the samples were then burned in a muffle furnace at 450°C for 6 hours. The samples were allowed to cool in the furnace before being weighed and the weight of the resulting ash was recorded. The difference between the ash weight and the dry weight was calculated to determine the AFDW of each sample. Host soluble protein content of each sample was determined via the Bradford method. Specifically, 10 µL of the coral fraction was pipetted into each of three wells of a 96-well plate (technical replicates), followed by the addition of 300 µL of Coomassie Plus Bradford reagent (Thermo Fisher Scientific) to each well. The plate was incubated for 10 minutes at room temperature followed by the collection of an absorbance reading at 595 nm on a spectrophotometer (BioTek PowerWave XS2). Each plate contained a set of internal standards with known concentrations of bovine serum albumin (0–2000 µg $mL^{-1}$), which were used to generate a second-degree polynomial standard curve relating absorbance (x) to protein concentration (y). The standard curve equation was used to calculate the protein concentrations of the samples. The concentration of lipids in each sample was determined via a modified method of (Dunn et al. 2012). First, ~45 mL of the coral fraction was lyophilized to produce dry tissue. A subsample of the lyophilized tissue was weighed (~90 mg), and 2 mL of chloroform-methanol (2:1) was added. The sample was then mixed using a homogenizer at 15000 rpm for 5 seconds, and an additional 2 mL of chloroform-methanol was used to rinse the homogenizer into the sample, and the tubes were then vigorously vortexed before being placed at 4°C for 2 hours in the dark. After incubation, tubes were again vortexed and the chilled homogenates were then passed through 0.22 µm syringe filters into new pre-weighed tubes. An additional 1 mL of chloroform-methanol was passed through the filters into each tube to ensure the full passage of sample material. Next, 1 mL of 0.1 M KCl was added to the samples, which were then vortexed and placed at 4°C for at least 1 hour in the dark until two phases formed. The aqueous (top) phase was discarded, and 5 mL of 50% methanol was added to the organic (bottom) phase. The samples were then placed at 4°C for 1 hour in the dark, followed by removal of the aqueous (top) phase. Washes with 50% methanol were repeated twice more, and the remaining organic phases were dried under a fume hood until the solvent had completely evaporated, at which point the lipid pellet was weighed. Coral host biomass, protein and lipid content were standardized to skeletal surface area.

The total antioxidant capacity of the coral homogenate was assessed using a kit from Cell Biolabs (STA-360) according to the manufacturer's instructions. First, 20 µL of the coral fraction was transferred to each of two wells of a 96-well plate (technical replicates), followed by the addition of 1x reaction buffer (20 µL) from the kit. An initial absorbance reading was collected with a spectrophotometer at 490 nm. Next, 50 µL of 1x copper ion reagent were added to each well, followed by incubation for 5 minutes with gentle shaking. Following shaking, 50 µL of 1x stop solution was added to each well, and a second absorbance reading was collected at 490 nm. Each plate contained a set of standards with known concentrations (0–0.1 mM) of uric acid. For analysis, the initial absorbance readings were subtracted from the final absorbance readings for each standard well, and the resulting values were used to generate a linear standard curve relating the absorbance (x) to the concentration of uric acid (y). The standard curve equation was used with the average change in absorbance across the two technical replicates for each sample to first determine the concentration of uric acid equivalents in the samples, which was then converted to a concentration of copper reducing equivalents (CRE) using the equivalence of 1 mM of uric acid to 2189 µM CREs. Finally, the CRE values were normalized to the amount of protein in each well, for a final value of TAC expressed as µM CRE mg protein$^{-1}$.

To assess the melanin content of the coral homogenate, a protocol was adapted from (Wall et al. 2021). First, 1.5 mL tubes were filled with approximately 20 mg of the dry coral fraction and the weight recorded. Next, 300 µL of 10 M NaOH was added to each tube, followed by vortexing for 20 seconds. Samples were left overnight at room temperature in the dark, then vortexed again for 10 seconds before being centrifuged at 7000 x g for 5 minutes. Following centrifugation, 100 µL of supernatant was transferred to each of two wells of a 96-well plate (technical replicates), which was then read for absorbance on a spectrophotometer at 490 nm. Each plate contained a set of internal standards with known concentrations (0–0.01 mg mL$^{-1}$) of synthetic melanin (M8631, Sigma Aldrich) in 10 M NaOH, which were used to generate a linear standard curve relating absorbance (x) to melanin concentration (y). The standard curve equation was then used to calculate the concentration of melanin in each sample, which was standardized to the original weight of the dry coral host tissue used in the assay.

A protocol was adapted from (Wall et al. 2021; Mydlarz and Palmer 2011; Fuess et al. 2018) to assess the content of prophenoloxidase (PPO) in each coral sample. First, 150 µL of the coral fraction was transferred to each of two wells of a 96-well plate (technical replicates). Next, 23 µL of 0.2 mg mL$^{-1}$ trypsin was added to each well, and the plate was incubated for 5 minutes at room temperature with shaking. Following incubation, 60 µL of 10 mM L-1,3-

dihydroxyphenylalanine (L-DOPA) was added to all wells, and the plate was again incubated at room temperature for 5 minutes with shaking. Next, the plate was read for absorbance on a spectrophotometer at 490 nm every minute for 15 minutes. The absorbance of each well at the start of the 15 minutes was subtracted from the absorbance of the same well at the end, and values were divided by 15 to determine the change in absorbance per minute, which were then averaged over the two technical replicates. Finally, these values were standardized to the amount of protein in each well for a final quantification of PPO activity expressed as the change in absorbance minute$^{-1}$ mg protein$^{-1}$.

Wax-dipping was used to determine calcium carbonate ($CaCO_3$) bulk density, where the skeleton was cleaned, dried to a constant mass and weighed, sealed with a coat of wax, dry weighed with the wax and then buoyant weighed in DI water at 20°C (Tambutté et al. 2015; K. T. Brown and Mello-Athayde 2022). The difference between dry weight (with wax) and buoyant weight (divided by the density of the DI water medium of 1 mg cm$^{-3}$) was calculated to determine the total volume enclosed. The dry skeletal mass (wax free) was then divided by the total volume enclosed to yield bulk density (g cm$^{-3}$).

*Acute heat stress experiment*

Ten individual colonies of each phenotype per species were sampled on September 1, 2022. Four fragments were collected from each individual colony (i.e., genetic clones) by hand using bone cutters, totalling 160 fragments. Following collection, corals were transported to the Hawai'i Institute of Marine Biology (HIMB) and placed in outdoor, flow-through seawater tanks under ambient temperatures (mean ± SD; 28.2 ± 0.1°C, n=2 tanks) until experimentation. Coral thermal tolerance experiments were initiated 72 hours after collection. The experimental heat stress assay followed the standardized temperature profile used to measure heat tolerance in corals (e.g., MMM, MMM+3°C, MMM+6°C, and MMM+9°C) (Voolstra et al. 2020). The experiment began on September 4, 2022, at 13:00 with a 3-hour ramp to respective treatment temperatures (27.5°C, 30.5°C, 33.5°C, 36.5°C), a 3-hour hold, and a 1-hour ramp down to MMM temperature (27.5°C). A fragment from each coral colony was suspended using fishing line and randomly placed into each treatment, so that all genotypes across species were present in each treatment. Temperatures were controlled using a custom computer-controlled system (Raspberry Pi). Temperatures were constantly monitored and manipulated by responding to probe measurements in real time and recorded using cross-calibrated temperature loggers (accuracy: ± 0.47°C at 25°C; resolution: ± 0.10°C at 25°C; HOBO UA-001-64, Onset Computer Corporation). Experiments were performed outdoors under natural photosynthetically active radiation (PAR), which ranged between 515–580 μmol m$^{-2}$ sec$^{-1}$ at midday (~12:00; Licor cosine sensor). At the end of the ramp and 1 h after sunset (~19:30), corals were assessed for dark-adapted photochemical yield ($F_v/F_m$) using a Diving-PAM (Walz GmbH) 5-mm diameter fiber-optic probe at a standardized distance (5 mm) above the coral tissue after $F_0$ stabilized. Three random spots (2–3 cm apart across the front and back of the nubbin) were measured on each fragment to obtain average measures of $F_v/F_m$. All readings with $F_0$ values that were less than 110 were removed to avoid any false detections (Marzonie et al. 2022). The following morning at 07:00 corals were photographed with a color standard (WDKK Waterproof Color Chart, DGK Color Tools) to assess coral color, a proxy for bleaching severity. Bleaching severity was determined visually (e.g., 0, 20, 40, 60, 80, 100% white) (Evensen et al. 2022) using the six gray standards as a reference (e.g., black= 0% bleached, white = 100% bleached) from two photographs (front and back of the fragment) to obtain an average color score (Fig. S4).

*Statistical analyses*

Graphical representations were produced using the package *ggplot2 (Wickham 2016)*. For colony-level bleaching severity, the interactive effects of phenotype (bleaching-susceptible, bleaching-resistant), coral species (*M. capitata*, *P. compressa*), season (summer: May, June, July, August, September, October; winter: November, December, January, February, March, April), and year (May 2015– April 2016, 2016–2017, 2017, 2019–2020, 2020–2021,2021–2022, 2022) were explored using a linear model. To assess differences in physiological parameters (total biomass, host soluble protein, lipids, $CaCO_3$ density, TAC, melanin content, PPO activity, CS, endosymbiont density, chlorophyll-*a* content) between phenotype, coral species and months post-heat stress (0, 10, 13, 17, 20, 24, 29, 35), linear mixed effects (lme) models were used, with genet (i.e., colony ID) included as a random effect.

Differences in temperature profiles of the experimental stress assays (four levels: ambient, ambient+3°C, ambient+6°C, ambient+9°C) were explored using a linear model. To determine how heat tolerance differed amongst coral species and phenotype, three-parameter log-logistic dose-response curves were fit to the average dark-adapted photochemical yield ($F_v/F_m$) across temperature treatments using the function dose-response model, drm, in the package *drc* (Ritz et al. 2015). From these dose-response curves, the effective temperature to induce a 50% loss in $F_v/F_m$ (effective dose 50; ED50) was calculated as the $F_v/F_m$ or color score from the model fit that is 50% of the initial value (Evensen et al. 2022).

Differences in coral multivariate phenotypes were analyzed separately for each coral species using permutational multivariate analysis of variance (PERMANOVA) and principal components analysis (PCA), with the fixed effects phenotype and months post-heat stress using the adonis and rda functions in the *vegan* package, respectively (Oksanen et al. 2013). Significant effects were followed by pairwise comparisons using the pairwiseAdonis2 function in the *pairwiseAdonis* package (Arbizu 2020). Resemblance matrices were obtained using Bray-Curtis dissimilarity and 9999 permutations.

**Results**

*Metabolic rates*

Maximum net photosynthesis ($P_{max}$) ($\chi^2 = 11.8$, $p = 0.008$) and light-enhanced dark respiration (LEDR) ($\chi^2 = 11.8$, $p = 0.008$) were significantly influenced by the interaction between coral species and time. Pairwise comparisons revealed that across all timepoints, $P_{max}$ and LEDR were greater in *P. compressa* than *M. capitata* ($p < 0.001$), and no significant differences were found between timepoints for either species ($p > 0.16$) (Fig. 3, Fig. S4). Photosynthesis to respiration (P:R) rates were significantly influenced by phenotype ($\chi^2 = 4.1$, $p = 0.04$), where bleaching-resistant corals trended towards higher P:R rates than bleaching-susceptible conspecifics (Fig. 3, Fig. S4).

*Immunity and antioxidant metrics*
Total antioxidant capacity (TAC) was significantly influenced by the interaction between coral species and time ($\chi^2 = 24.2$, $p = 0.001$). For *M. capitata*, TAC was highest in October 2021 and significantly greater than the same time during the previous year (November 2020), as well as in comparison to the marine heatwave (October 2019) ($p < 0.003$) (Fig. 3g, Fig. S4). Similarly for *P. compressa*, TAC was significantly greater in November 2020 and October 2021 when compared to the 2019 marine heatwave ($p < 0.05$) (Fig. 3g, Fig. S4).

Melanin content was significantly influenced by the interaction between coral species and time ($\chi^2 = 17.1$, $p = 0.009$). Melanin concentrations were not measured during the 2019 marine heatwave. Across the time series, *P. compressa* had higher melanin concentrations than *M. capitata* ($p < 0.002$) (Fig. 3h, Fig. S4). For *M. capitata*, melanin concentrations did not differ between time points ($p > 0.27$), whereas for *P. compressa*, seasonal patterns were observed with concentrations generally greatest between March–June, yet also elevated in October 2021 (Fig. 3h).

Prophenoloxidase (PPO) activity was significantly influenced by the three-way interaction between coral species, phenotype and time ($\chi^2 = 18.2$, $p = 0.006$). Pairwise comparisons revealed that bleaching-susceptible *P. compressa* had lower PPO activity than bleaching-resistant corals 13 months post-heat stress and bleaching-susceptible *M. capitata* had lower PPO activity than bleaching-resistant conspecifics 17 months post-heat stress (Fig. 3i, Fig. S4). For several time points (13, 24, and 29 months post-heat stress), *P. compressa* had greater PPO activity than *M. capitata*, particularly bleaching-resistant comparisons ($p < 0.04$). PPO activity was significantly lower in August–September (10 months and 35 months post-heat stress) than all other time points regardless of coral species or phenotype ($p < 0.05$) (Fig. 3i, Fig. S4).

*Short-term heat stress experiment*
Color score, as a proxy for bleaching severity, was significantly influenced by the interaction between species and phenotype ($X^2 = 4.1$, $p = 0.04$) as well as individual effect of temperature ($X^2 = 134.1$, $p < 0.0001$). There were no significant differences in $F_v/F_m$ between *M. capitata* and *P. compressa* at ambient ($p = 0.09$), +3°C ($p = 0.06$), or +6°C ($p = 0.08$), yet in the hottest treatment (+9°C), *P. compressa* $F_v/F_m$ was significantly greater than *M. capitata* ($p < 0.0001$) (Fig. 5). Pairwise comparison revealed no differences in bleaching severity between phenotypes of *P. compressa* ($p = 0.85$), yet for *M. capitata*, susceptible corals were significantly paler than resistant conspecific corals ($p < 0.0001$) (Fig. S6b). Regardless of species, bleaching severity increased across all treatments ($p <0.0001$), with the exception of ambient and +3°C ($p = 0.65$). Interestingly, heat tolerance (ED50) determined through color score was between 1.9 – 2.8°C lower than that determined via photochemical yield and there was only 0.4°C difference in the color score ED50 between the least and most heat tolerant species.

**Supplementary Figures and Tables**

**Table S1. Temperature metadata.** Abbreviations: NOAA = National Oceanic and Atmospheric Administration, PMEL = Pacific Marine and Environmental Laboratory

<table>
<tr><th>Contributor</th><th>Reef</th><th>Coordinates</th><th>Date range</th><th>Sensor type</th><th>Sampling interval</th><th>Depth (m)</th></tr>
<tr><td>NOAA - PMEL</td><td>CRIMP2</td><td>21.458, -157.798</td><td>2013-11-28 to 2020-03-30*</td><td>SBE 37 SMP[1]</td><td>3 hours</td><td>0.7</td></tr>
<tr><td>HIMB weather station</td><td>Reef 1</td><td>21.433, -157.7863</td><td>2014-01-01 to 2022-11-22*</td><td>BetaTherm 100K6A1IA Thermistor[2]</td><td>1 hour</td><td>1.0</td></tr>
<tr><td rowspan="2">Division of Aquatic Resources (DAR)</td><td rowspan="2">Reef 12</td><td>21.45096, -157.79723</td><td rowspan="2">2015-06-19 to 2016-01-21</td><td rowspan="2">Onset Pro v2[3]</td><td rowspan="2">15 minutes</td><td rowspan="2">DH 1.5</td></tr>
<tr><td>21.45090, -157.79729</td></tr>
<tr><td>Gates Lab (Katie Barott)</td><td>Reef 13</td><td>21.4516, -157.7966</td><td>2016-07-06 to 2017-04-18</td><td>SBE Sea-pHOx[1]</td><td>15 minutes</td><td>2.0</td></tr>
<tr><td rowspan="4">Coral Reef Resilience Lab (Carlo Caruso, Crawford Drury, Mariana Rocha de Souza)</td><td rowspan="4">Reef 13</td><td>21.4518, -157.7956</td><td rowspan="2">2019-09-27 to 2021-08-26</td><td rowspan="4">Onset Pro v2[3]</td><td rowspan="4">10 minutes</td><td>1.2</td></tr>
<tr><td>21.4509, -157.7954</td><td>1.7</td></tr>
<tr><td>21.4518, -157.7956</td><td rowspan="2">2022-02-26 to 2022-11-28</td><td>1.2</td></tr>
<tr><td>21.4509, -157.7954</td><td>1.7</td></tr>
<tr><td rowspan="3">Barott Lab (Kristen Brown and Katie Barott)</td><td rowspan="3">Reef 13</td><td>21.4516, -157.7965</td><td rowspan="3">2022-09-02 to 2023-04-13</td><td rowspan="3">Onset Pro V2[3]</td><td rowspan="3">30 minutes</td><td>1.0</td></tr>
<tr><td>21.4507, -157.7962</td><td>1.0</td></tr>
<tr><td>21.4517, -157.7967</td><td>2.5</td></tr>
</table>

*Incomplete record.

[1]Accuracy: ±0.002°C; resolution: 0.0001°C at 25°C

[2]Tolerance: ±0.2°C; Steinhart-Hart equation error: ≤ ±0.01°C at 25°C

[3]Accuracy: ±0.21°C; resolution: 0.02°C at 25°C

**Table S2. Comparison of maximum cumulative degree heating weeks from 2014–2022.** Light gray shading indicates years with recognized marine heatwaves.

| Maximum cumulative degree heating weeks | | | | | |
|---|---|---|---|---|---|
| Location | Kāneʻohe Bay^ | | | | Main Hawaiian Islands* |
| Temperature dataset | Reef 13 | Reef 1 (Moku o Loʻe) | Reef 13 | Reef 1 (Moku o Loʻe) | NOAA (SST) |
| Year | 24 hour mean; Local MMM (27.3°C) | | 24 hour mean; Regional MMM (27.0°C) | | Nighttime mean; Regional MMM (27.0°C) |
| 2014 | 7.3 | 5.2 | 10.2 | 9.0 | 4.5 |
| 2015 | 8.8 | 7.2 | 14.6 | 11.2 | 12.5 |
| 2016 | 0 | 0 | 0.2 | 0.3 | 0 |
| 2017 | 0.6 | 0 | 3.2 | 2.0 | 1.3 |
| 2018 | 0.8 | 0.6 | 2.0 | 1.5 | 0.15 |
| 2019 | 5.1 | 10.2 | 10.2 | 15.3 | 13.6 |
| 2020 | 0.1 | 0.2 | 3.1 | 1.0 | 0.5 |
| 2021 | 0 | 0.2 | 0.7 | 0.7 | 0 |
| 2022 | 0.2 | 0.6 | 2.9 | 2.1 | 0 |

^Kāneʻohe Bay data are from *in situ* temperature loggers (depth 0.7 – 2.7 m; as described in Table S1).

*Main Hawaiian Islands data are from NOAA satellite-derived sea surface temperature (SST).

**Table S3.** Endosymbiont community composition of *Montipora capitata* colonies sampled for this study. Symbiont percentages reflect quantitative PCR results from colony samples collected July 9, 2019 (Dilworth et al. 2021).

| Colony ID | Bleaching phenotype | *Durusdinium glynnii* (%) | *Cladocopium* 'C31' (%) |
|---|---|---|---|
| 3 | Susceptible | 0 | 100 |
| 4 | Resistant | 83 | 17 |
| 11 | Susceptible | 41 | 59 |
| 12 | Resistant | 98 | 2 |
| 19 | Susceptible | 0 | 100 |
| 20 | Resistant | 98 | 2 |
| 201 | Susceptible | 0 | 100 |
| 202 | Resistant | 92 | 8 |
| 203 | Susceptible | 0 | 100 |
| 204 | Resistant | 96 | 4 |
| 209 | Susceptible | 0 | 100 |
| 210 | Resistant | 64 | 36 |
| 211 | Susceptible | 0 | 100 |
| 212 | Resistant | 98 | 2 |
| 213 | Susceptible | 1 | 99 |
| 214 | Resistant | 98 | 2 |
| 219 | Susceptible | 0 | 100 |
| 220 | Resistant | 98 | 2 |
| 221 | Susceptible | 0 | 100 |
| 222 | Resistant | 97 | 3 |

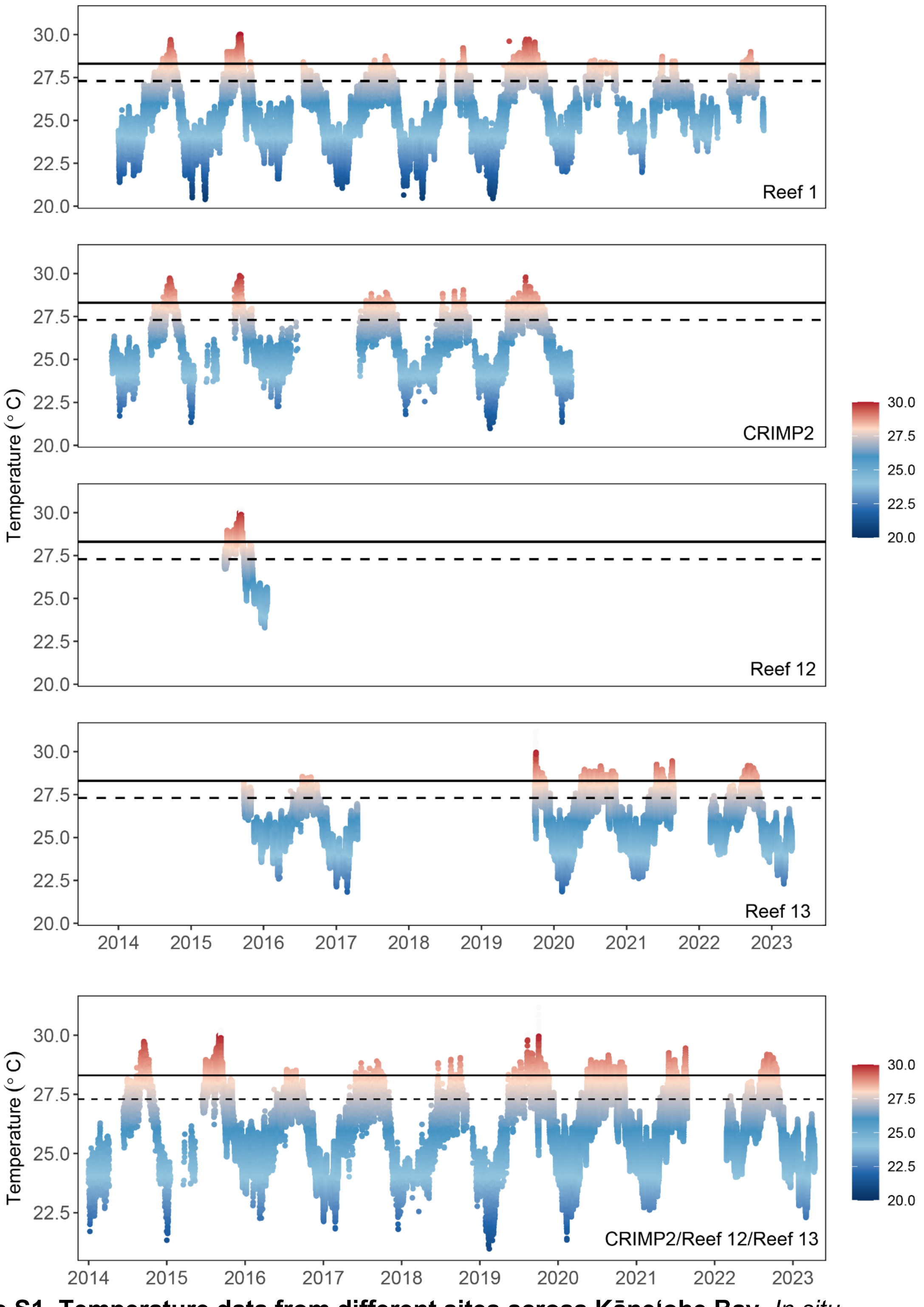


**Figure S1. Temperature data from different sites across Kāne'ohe Bay.** *In situ* temperatures were recorded from January 2014 – April 2023 at a depth of 0.7–2.7 m in Kāne'ohe Bay. Points indicate hourly measurements. Dashed horizontal line indicates the Kāne'ohe Bay's climatological maximum monthly mean (MMM; 27.3°C) and solid horizontal line indicates the local coral bleaching threshold (MMM+1°C; 28.3°C).

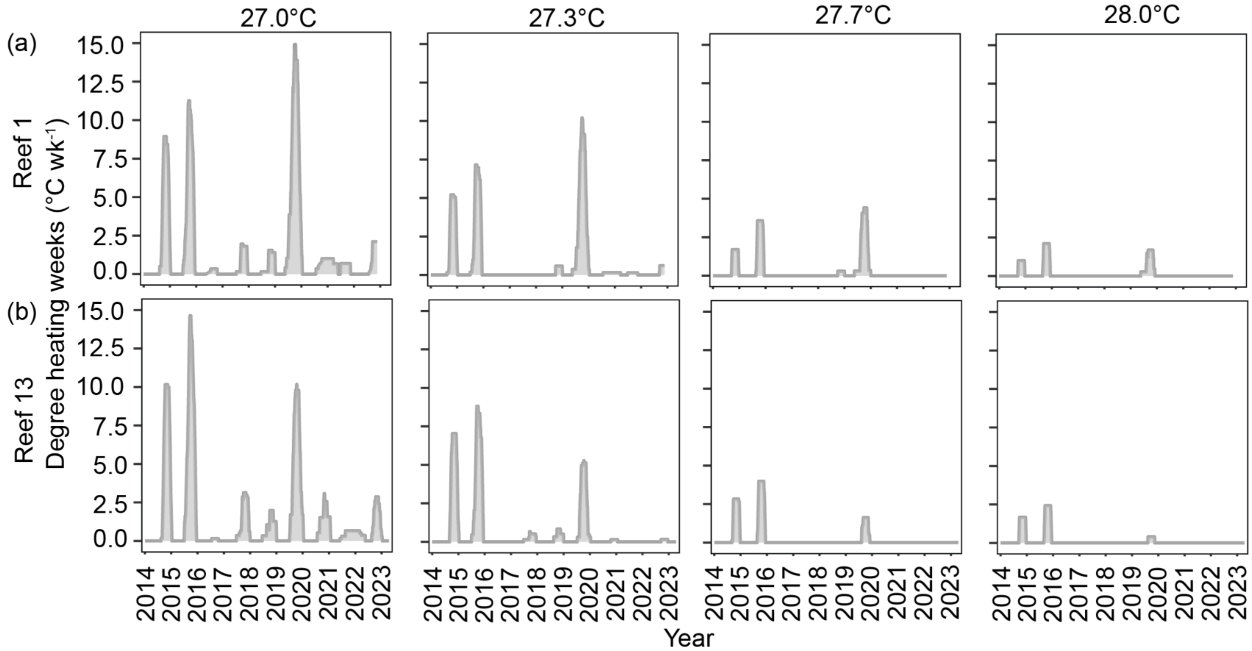


**Figure S2. Comparison of degree heating weeks across Kāne'ohe Bay.** (a) Degree heating week profiles at at Patch Reef 1 (Moku o Lo'e) calculated using a maximum monthly mean (MMM) of 27.0°C (current regional MMM used by NOAA for the Main Hawaiian Islands) and a range of MMM used for Kāne'ohe Bay: 27.0°C, climatic MMM from 1960s (Jokiel and Coles 1977); 27.3°C, climatic MMM from 1990s; 27.7°C, used by (Wall et al. 2021; Jury and Toonen 2019), and 28.0°C, used by (Bahr, Jokiel, and Rodgers 2015; Jury and Toonen 2019; Innis et al. 2021; Matsuda et al. 2020). (b) Degree heating week profiles calculated using a MMM of 27.0°C, 27.3°C, 27.7°C, and 28.0°C at Patch Reef 13.

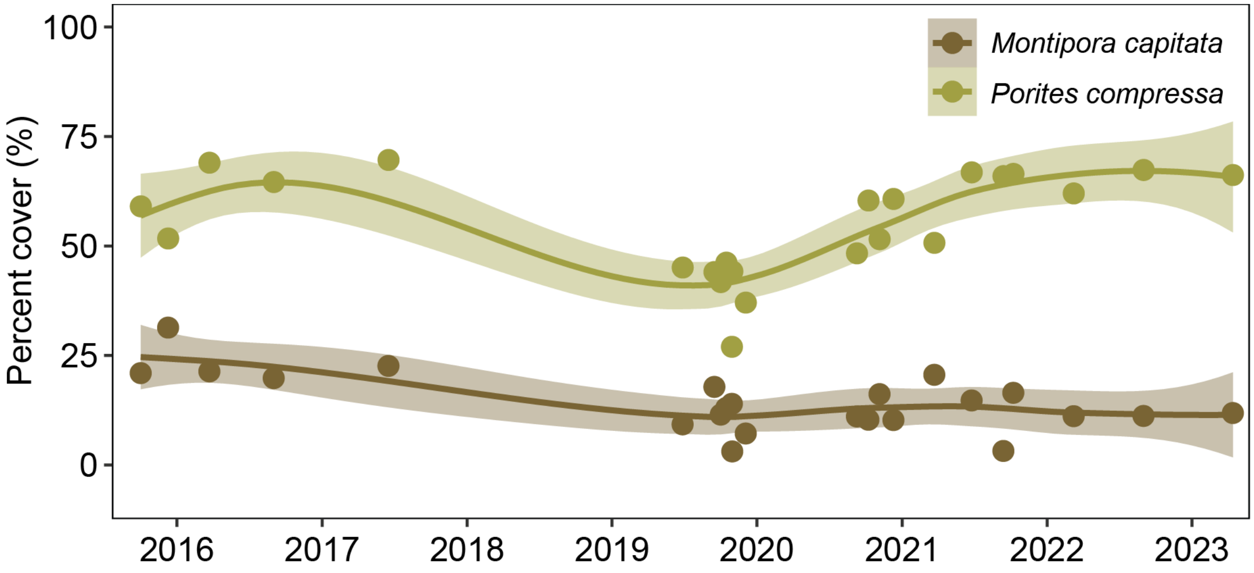


**Figure S3.** Hard coral cover of species of interest (*Montipora capitata* and *Porites compressa*) at patch reef 13 from September 2015–September 2022. Points represent mean percent cover from benthic surveys (n = 2–4), and lines and ribbons represent smoothed conditional means ± 95% CI.

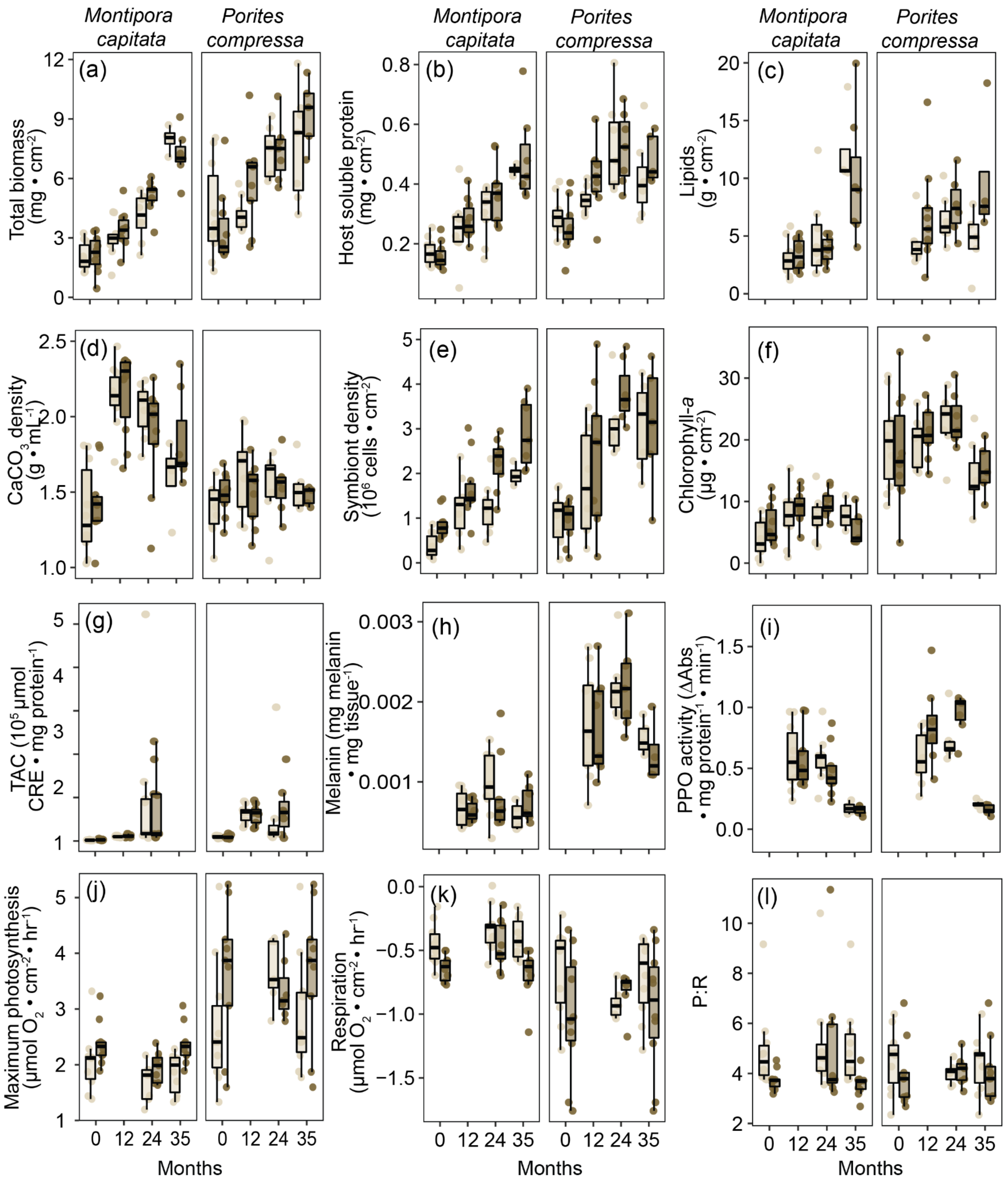


**Figure S4. Physiological traits of bleaching-resistant and bleaching-susceptible corals during peak annual temperatures in the years following the 2019 heatwave.** Months indicate time post-heat stress, where ‘0’ represents during the 2019 marine heatwave. (a) Host tissue biomass (ash-free dry weight), (b) host soluble protein density, (c) host lipids, (d) calcium carbonate ($CaCO_3$) density, (e) endosymbiont cell density, (f) chlorophyll *a* concentration, (g) host total antioxidant capacity (TAC), (h) host melanin content, (i) host prophenoloxidase (PPO)

activity, (j) maximum photosynthesis, (k) light-enhanced dark respiration, and (l) photosynthesis to respiration ratios (P:R) for bleaching-susceptible and bleaching-resistant *Monitpora capitata* and *Porites compressa*. Boxplots display the minimum, 25th percentile, median, 75th percentile, and maximum, where points indicate individual measures for coral genets (n = 7–10).

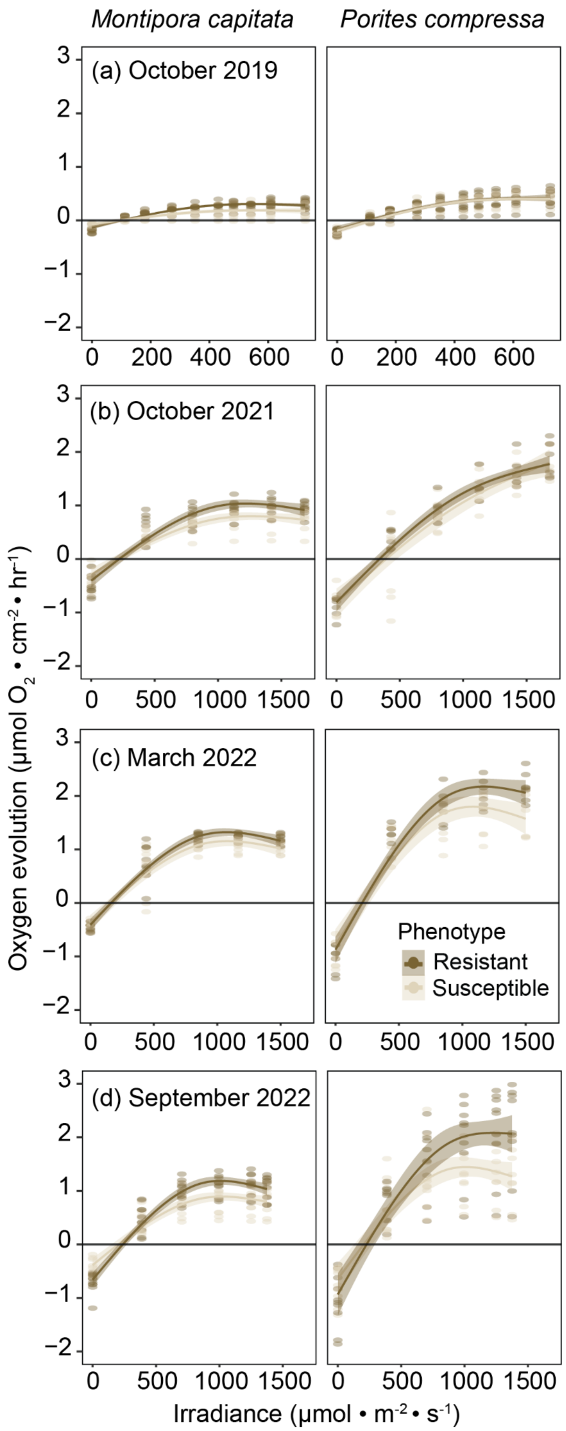


**Figure S5.** Photosynthesis–irradiance curves of bleaching-susceptible and bleaching-resistant *Monitipora capitata* and *Porites compressa* in (a) October 2019 (during the heatwave; note different x-axis scale), (b) October 2021, (c) March 2022, and (d) September 2022.

Bahr, Keisha D., Paul L. Jokiel, and Kuʻulei S. Rodgers. 2015. "The 2014 Coral Bleaching and Freshwater Flood Events in Kāneʻohe Bay, Hawaiʻi." *PeerJ* 3 (August): e1136.

Beijbom, Oscar, Peter J. Edmunds, Chris Roelfsema, Jennifer Smith, David I. Kline, Benjamin P. Neal, Matthew J. Dunlap, et al. 2015. "Towards Automated Annotation of Benthic Survey Images: Variability of Human Experts and Operational Modes of Automation." *PloS One* 10 (7): e0130312.

Brown, Kristen T., Gal Eyal, Sophie G. Dove, and Katie L. Barott. 2022. "Fine-Scale Heterogeneity Reveals Disproportionate Thermal Stress and Coral Mortality in Thermally Variable Reef Habitats during a Marine Heatwave." *Coral Reefs* , November, 1–12.

Brown, K. T., and M. A. Mello-Athayde. 2022. "Environmental Memory Gained from Exposure to Extreme pCO2 Variability Promotes Coral Cellular Acid–base Homeostasis." *Of the Royal …*. https://royalsocietypublishing.org/doi/abs/10.1098/rspb.2022.0941.

Dilworth, Jenna, Carlo Caruso, Valerie A. Kahkejian, Andrew C. Baker, and Crawford Drury. 2021. "Host Genotype and Stable Differences in Algal Symbiont Communities Explain Patterns of Thermal Stress Response of Montipora Capitata Following Thermal Pre-Exposure and across Multiple Bleaching Events." *Coral Reefs* 40 (1): 151–63.

Dunn, Simon R., Michael C. Thomas, Geoffrey W. Nette, and Sophie G. Dove. 2012. "A Lipidomic Approach to Understanding Free Fatty Acid Lipogenesis Derived from Dissolved Inorganic Carbon within Cnidarian-Dinoflagellate Symbiosis." *PloS One* 7 (10): e46801.

Edmunds, P. J., and P. Spencer Davies. 1988. "Post-Illumination Stimulation of Respiration Rate in the Coral Porites Porites." *Coral Reefs* 7 (1): 7–9.

Evensen, Nicolas R., Christian R. Voolstra, Maoz Fine, Gabriela Perna, Carol Buitrago-López, Anny Cárdenas, Guilhem Banc-Prandi, Katherine Rowe, and Daniel J. Barshis. 2022. "Empirically Derived Thermal Thresholds of Four Coral Species along the Red Sea Using a Portable and Standardized Experimental Approach." *Coral Reefs* 41 (2): 239–52.

Fuess, Lauren E., Whitney T. Mann, Lea R. Jinks, Vanessa Brinkhuis, and Laura D. Mydlarz. 2018. "Transcriptional Analyses Provide New Insight into the Late-Stage Immune Response of a Diseased Caribbean Coral." *Royal Society Open Science* 5 (5): 172062.

Holmes, Glen. 2008. "Estimating Three-Dimensional Surface Areas on Coral Reefs." *Journal of Experimental Marine Biology and Ecology* 365 (1): 67–73.

Innis, Teegan, Luella Allen-Waller, Kristen T. Brown, Wesley Sparagon, Christopher Carlson, Elisa Kruse, Ariana S. Huffmyer, Craig E. Nelson, Hollie M. Putnam, and Katie L. Barott. 2021. "Marine Heatwaves Depress Metabolic Activity and Impair Cellular Acid-Base Homeostasis in Reef-Building Corals regardless of Bleaching Susceptibility." *Global Change Biology* 27 (12): 2728–43.

Jeffrey, S. W., and G. F. Humphrey. 1975. "New Spectrophotometric Equations for Determining Chlorophylls A, B, c1 and c2 in Higher Plants, Algae and Natural Phytoplankton." *Biochemie Und Physiologie Der Pflanzen: BPP* 167 (2): 191–94.

Jokiel, P. L., and S. L. Coles. 1977. "Effects of Temperature on the Mortality and Growth of Hawaiian Reef Corals." *Marine Biology* 43 (3): 201–8.

Jury, Christopher P., and Robert J. Toonen. 2019. "Adaptive Responses and Local Stressor Mitigation Drive Coral Resilience in Warmer, More Acidic Oceans." *Proceedings of the Royal Society B: Biological Sciences* 286 (1902): 20190614.

Marzonie, Magena R., Line K. Bay, David G. Bourne, Andrew S. Hoey, Samuel Matthews, Josephine J. V. Nielsen, and Hugo B. Harrison. 2022. "The Effects of Marine Heatwaves on Acute Heat Tolerance in Corals." *Global Change Biology*, October. https://doi.org/10.1111/gcb.16473.

Matsuda, Shayle, Ariana Huffmyer, Elizabeth A. Lenz, Jen Davidson, Joshua Hancock, Ariana Przybylowski, Teegan Innis, Ruth D. Gates, and Katie L. Barott. 2020. “Coral Bleaching Susceptibility Is Predictive of Subsequent Mortality within but Not between Coral Species.” *Frontiers in Ecology and Evolution* 8 (178): 1–14.

Mydlarz, Laura D., and Caroline V. Palmer. 2011. “The Presence of Multiple Phenoloxidases in Caribbean Reef-Building Corals.” *Comparative Biochemistry and Physiology. Part A, Molecular & Integrative Physiology* 159 (4): 372–78.

Oksanen, Jari, F. Guillaume Blanchet, Roeland Kindt, Pierre Legendre, Peter R. Minchin, R. B. O’hara, Gavin L. Simpson, Peter Solymos, M. Henry H. Stevens, and Helene Wagner. 2013. “Package ‘vegan.’” *Community Ecology Package, Version* 2 (9): 1–295.

Platt, T., C. L. Gallegos, and W. G. Harrison. 1981. “Photoinhibition of Photosynthesis in Natural Assemblages of Marine Phytoplankton.” repositorio.imarpe.gob.pe. 1981. https://repositorio.imarpe.gob.pe/bitstream/20.500.12958/1357/1/BOL%20EXTR.%20Investigaci%C3%B3n%20....-18.pdf.

Ritz, Christian, Florent Baty, Jens C. Streibig, and Daniel Gerhard. 2015. “Dose-Response Analysis Using R.” *PloS One* 10 (12): e0146021.

Rodgers, K. S., P. L. Jokiel, and Western Weather Group, Inc. 2005. “HIMB Weather Station: Moku O Loe (Coconut Island), Oahu, Hawaii. [1992-2002] Distributed by the Pacific Islands Ocean Observing System (PacIOOS).” http://pacioos.org/metadata/aws_himb.html.

Stimson, John, and Robert A. Kinzie. 1991. “The Temporal Pattern and Rate of Release of Zooxanthellae from the Reef Coral Pocillopora Damicornis (Linnaeus) under Nitrogen-Enrichment and Control Conditions.” *Journal of Experimental Marine Biology and Ecology* 153 (1): 63–74.

Tambutté, E., A. A. Venn, Michael Holcomb, N. Segonds, N. Techer, D. Zoccola, D. Allemand, and S. Tambutté. 2015. “Morphological Plasticity of the Coral Skeleton under CO 2-Driven Seawater Acidification.” *Nature Communications* 6 (1): 1–9.

Voolstra, Christian R., Carol Buitrago-López, Gabriela Perna, Anny Cárdenas, Benjamin C. C. Hume, Nils Rädecker, and Daniel J. Barshis. 2020. “Standardized Short-Term Acute Heat Stress Assays Resolve Historical Differences in Coral Thermotolerance across Microhabitat Reef Sites.” *Global Change Biology*, June. https://doi.org/10.1111/gcb.15148.

Wall, Christopher B., Contessa A. Ricci, Alexandra D. Wen, Bren E. Ledbetter, Delania E. Klinger, Laura D. Mydlarz, Ruth D. Gates, and Hollie M. Putnam. 2021. “Shifting Baselines: Physiological Legacies Contribute to the Response of Reef Corals to Frequent Heatwaves.” *Functional Ecology* 35 (6): 1366–78.

Wickham, Hadley. 2016. *ggplot2: Elegant Graphics for Data Analysis*. Springer.
